# Assessing bnAb potency in the context of HIV-1 Envelope conformational plasticity

**DOI:** 10.1101/2024.09.13.612684

**Authors:** Caio Foulkes, Nikolas Friedrich, Branislav Ivan, Emanuel Stiegeler, Carsten Magnus, Daniel Schmidt, Umut Karakus, Jacqueline Weber, Huldrych F. Günthard, Chloé Pasin, Peter Rusert, Alexandra Trkola

## Abstract

The ability of broadly neutralizing antibodies (bnAbs) to interact with the closed, pre-fusion HIV-1 envelope (Env) trimer distinguishes them from weakly neutralizing antibodies (weak-nAbs) that depend on trimer opening to bind. Comparative analysis of neutralization data from the CATNAP database revealed a nuanced relationship between bnAb activity and Env conformational plasticity, with substantial epitope-specific variation of bnAb potency ranging from increased to decreased activity against open, neutralization-sensitive Env. To systematically investigate the impact of Env conformational dynamics on bnAb potency we screened 126 JR-CSF point mutants for generalized neutralization sensitivity to weak-nAbs and plasma from people with chronic HIV-1 infection. 23 mutations at highly conserved sites resulted in neutralization phenotype with high Tier 1 sensitivity, which was associated with destabilization of the closed, prefusion conformation. Including 19 of these mutants into a Sensitivity Env mutant panel (SENSE-19), we classified bnAbs according to potency variations in response to trimer opening. To verify that these sensitivity patterns are independent of the in vitro assay system, replication-competent SENSE-19 mutant viruses were tested on primary CD4 T cells. While loss of potency on SENSE-19 was registered for bnAbs recognizing quaternary epitopes on pre-triggered Env, structural destabilization benefitted MPER bnAbs and other inhibitors known to have post-CD4 attachment neutralization activity. Importantly, for certain bnAbs targeting CD4bs, V3-glycan and interface epitopes, particularly low potency variation was noted, suggesting that Env conformational tolerance can be achieved but is not the rule. In summary, SENSE-19 screens revealed distinct Env flexibility tolerance levels between bnAb types that provide mechanistic insights in their function and broaden current neutralization breadth assessments.

**Author summary:** Consistently high potency and neutralizing breadth against genetically divergent strains circulating worldwide are central to the applicability of HIV-1 broadly neutralizing antibodies (bnAbs) for prevention and therapy. The trimeric Envelope protein complex on these viruses is the target for nAbs and because of its inherent flexibility can be presented in different shapes. However, the activity of nAbs depends to different degree on the presentation of certain shapes, limiting their potency and breadth. Here we assemble a virus panel that can be used to estimate this dependence, enabling the identification of bnAbs tolerating the presentation of different shapes.

## Introduction

Broadly neutralizing antibodies (bnAbs) are central to ongoing efforts to develop effective HIV-1 vaccines and therapeutics [1–3]. The principal capacity of bnAbs to counter the extraordinary diversity of HIV-1 strains is compelling but bnAb breadth and potency need to be at highest possible level to register in in vivo efficacy as recently demonstrated in animal models [4, 5] and by the Antibody Mediated Protection (AMP) trial in humans [6].

Activity of neutralizing antibodies (nAbs), including bnAbs, is steered by the accessibility of their epitopes on the intact HIV-1 envelope protein (Env) trimer [4, 5, 7–13]. Epitope specific alterations in amino acid composition or glycosylation conferring resistance to a distinct nAb class may often have no consequence on other nAb types. Generalized effects on neutralization sensitivity affecting multiple nAb epitopes are, however, observed when modulations in Env conformational stability occur [14–17]. Conformational flexibility and plasticity are inherent features of the HIV-1 Env trimer required for its function in entry [9, 17–20]. This inherent structural flexibility of Env, reflected also in its genetic variability, provides means for evasion by mutating and conformational masking of neutralization sensitive epitopes [11, 21–24]. During host cell entry the trimeric Env protein undergoes a series of conformational changes which are driven by sequential engagement of the primary receptor CD4 and a co-receptor ultimately leading to fusion of the virus and host cell membranes. The structural changes in the Env trimer upon CD4 binding are characterized by the transition from a “closed” to downstream “open” conformations exposing the co-receptor binding site as well as additional neutralization sensitive epitopes [8, 9, 18, 20, 25].

Typically, gross increases in Env neutralization sensitivity coincide with altered entry kinetics [15, 16, 26–28] and high Env conformational flexibility [14, 15, 17, 25] [14, 15, 17]. This flexibility provides access to multiple nAb epitopes and corresponding viruses are classified as Tier 1 strains in the neutralization sensitivity ranking of viral strains [29]. The Tier-system [29], a relative measure for the sensitivity of individual HIV-1 strains to Ab-mediated neutralization, is based on the resistance of HIV-1 strains to polyclonal Abs in plasma from people with HIV-1 (PWH) derived in chronic HIV-1 infection. Lab-adapted and highly sensitive primary HIV-1 strains are categorized as Tier 1A strains. Tier 1B, Tier 2 and Tier 3 strains are increasingly resistant to plasma neutralization, with Tier 2 comprising the majority of strains circulating worldwide [29].

The two main quality measures of bnAbs are their neutralization breadth against genetically divergent HIV-1 strains including different subtypes and the potency at which these strains are neutralized. In an effort to define bnAbs with the highest breadth and potency, their neutralization activity against >100, in some cases close to 200 virus strains from different clades is assessed in Env pseudovirus neutralization assays [30–32]. Algorithms for predicting bnAb epitopes and delineating neutralizing reactivity in polyclonal PWH plasma have been developed based on the wealth of information available from detailed analysis of bnAb neutralizing activity against these divergent strains [33, 34]. While the careful analysis of lead bnAbs on such large virus panels is highly informative, initial screening of patient plasmas and monoclonal antibodies (mAbs) must rely on smaller virus panels to enable higher sample throughput in determining neutralization breadth [35, 36]. Here we report on an alternative approach to identify and characterize lead bnAbs employing single site Env mutants with affirmed effects on trimer opening. While bnAbs show relative constant potency across wildtype and mutant Envs, this is not the case for weak-nAbs. As we show, systematic evaluation of the tolerance of bnAbs to trimer opening provides a measure to classify bnAbs according to their breadth and potency.

## Results

### Assessing bnAb potency in the context of HIV-1 Env conformational plasticity

bnAbs, per definition, inhibit genetically divergent virus strains and are known to retain high activity against Tier-2 viruses known to largely maintain a closed prefusion Env conformation[7, 17, 20, 37, 38]. While effects of modulating Env stability on neutralization activity have been described for some bnAbs [15, 16, 39, 40], a refined understanding of how Env conformational plasticity affects neutralization efficacy across bnAb types remains important. To compare bnAb activity over the full spectrum of highly flexible to less flexible Envs, we used Tier-based neutralization data available in the CATNAP database (http://hiv.lanl.gov/catnap)[41] (Fig 1). We selected 18 prototypic bnAbs covering the five major bnAb epitopes for this analysis (CD4-binding site (CD4bs) bnAbs NIH45-46, PGV04, 3BNC117, and VRC01; V2-glycan bnAbs PG9, PG16, PGDM1400, and PGT145; the gp120-gp41 interface bnAb PGT151; the V3-glycan bnAbs PGT121, PGT 128, PGT130, PGT135, BG18 and 2G12; and bnAbs against the membrane proximal external region (MPER) in gp41: 4E10, 10E8, and 2F5). Several weak-nAbs that represent Ab specificities prevalent at high titers in HIV-1 infection were chosen for comparison [42, 43]. These included nAbs to the variable region 3 (V3-crown: 447-52D), the CD4-induced epitope (CD4i: 17b) and agents directed to the CD4bs (nAbs 1F7 and b12, and the CD4 receptor analogue CD4-IgG2). Access to these epitopes requires either partial or full, spontaneous or receptor-induced conformational opening of the trimeric Env complex [9, 13, 44]. Geometric mean IC50 values stratified by neutralization Tier showed the expected pattern with weak-nAb activity being almost exclusively restricted to Tier 1 strains (Fig 1). This is in line with the more tightly closed Env prefusion conformation of higher tiered Envs disfavoring antibody attack by increased conformational shielding and/or altering entry kinetics[14–17, 25, 28].

**Fig 1.**
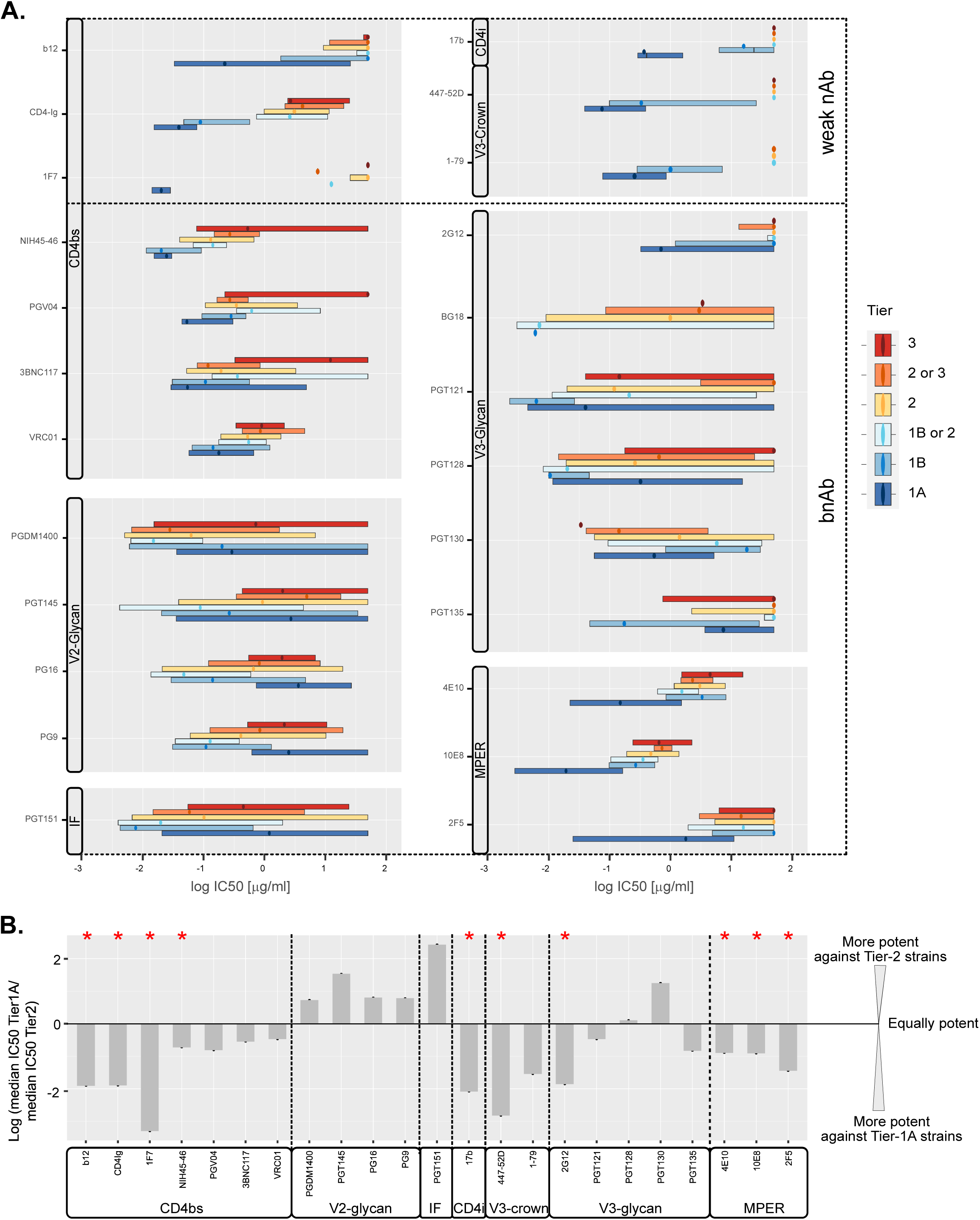
Ab-specific variations in neutralization potency against HIV-1 strains assigned to different neutralization tiers. All IC50 data available from the Los Alamos National Laboratory database (http://hiv.lanl.gov/catnap; as of February 2023) was downloaded for the antibodies listed (grouped by epitope as indicated) against all HIV-1 strains that had been attributed to a neutralization tier (tier 1A: most sensitive; tier 3: most resistant, total of 530 viruses). **A.** Distribution of IC50 values for each antibody and for HIV-1 strains of different tiers (dots: median, boxplot: values within the interquartile range; whiskers indicating minimum and maximum values were omitted for clarity). **B.** Summary of differences between median potencies of Abs against Tier 1A and Tier 2 strains. Significant differences (p<0.05, Mann-Whitney test) are indicated by a red star.

Interestingly, side-by-side comparison of the inhibitory activity of different bnAb classes against tiered viruses revealed that bnAbs also benefit from some degree of Env trimer opening. A gradual decrease in potency with increase in Tier-level was evident for most bnAbs (Fig 1A). CD4bs and MPER directed bnAbs inhibited Tier 1A viruses most potently. V2-glycan bnAbs and the interface directed bnAb PGT151 showed a multi-phase activity pattern with Tier 1A and Tier 3 strains recording the lowest neutralization sensitivity and peak potency against Tier1B or Tier 1B/2 classified strains. Patterns of V3-glycan bnAbs were diverse, but most of the tested V3-glycan bnAbs showed highest potency against either Tier 1A or Tier 1B viruses. While differences in median IC50 between Tier 2 and Tier 1A for bnAbs were in most cases within 0.5-1 orders of magnitude (Fig 1B), the differences for weak-nAbs remained the most pronounced (>1.5 orders of magnitude) leading in several cases to a switch from no activity to potent neutralization which is prototypical for mAbs against CD4i and V3-crown epitopes [11, 16, 45].

### Harnessing perturbations in Env conformation to classify bnAb activity

The higher activity of most bnAbs against lower-tier viruses suggests a general effect of relaxing Env conformational stability on bnAb potency. We therefore hypothesized that patterns of neutralizing activity based on Env stability could be used to classify nAbs. To explore this in a controlled setting we utilized a panel of single-site gp120 mutants of the subtype B Tier 2 strain JR-CSF covering 125 of the 503 gp120 amino acid positions [31, 46, 47] (Fig 2 and S1 Fig). To select mutants with a generally enhanced neutralization sensitivity, JR-CSF mutant pseudoviruses were screened against plasma from people with chronic, untreated HIV-1 infection (N=18; Table S1), the CD4 receptor analogue CD4-IgG2 and a range of weak-nAbs (CD4bs: b6 and b12, V3-crown: 1-79, 447-52D).

**Fig 2.**
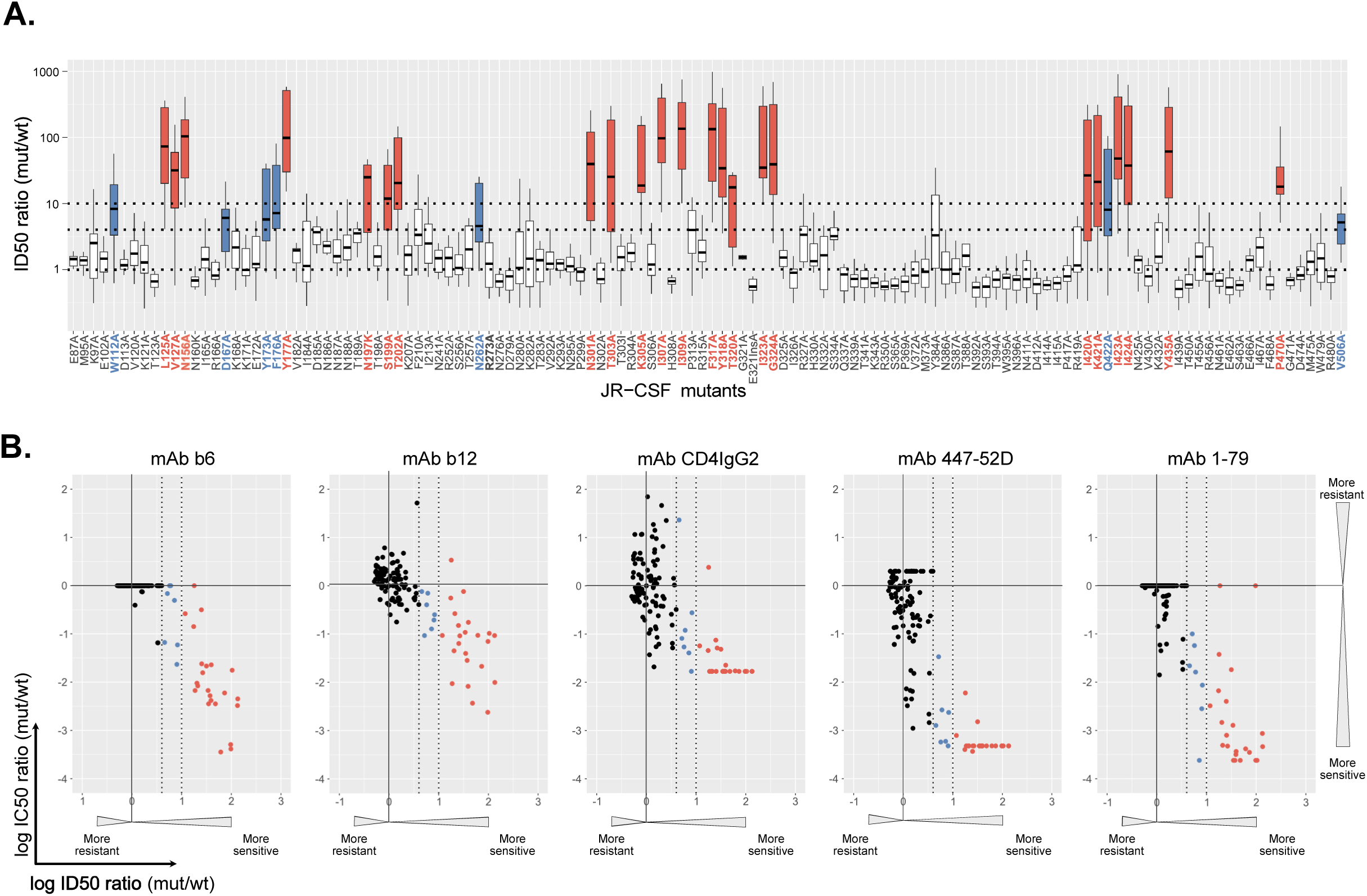
Characterizing the generalized neutralization sensitivity of JR-CSF envelope point mutants. The neutralization sensitivity of 126 JR-CSF envelope (Env) mutants was determined in a TZM-bl based pseudovirus inhibition assay. **A.** Each mutant virus was titrated with eleven plasma samples from HIV-1 chronically infected individuals (nine subtype B infected, two infected with non-B subtypes). All eleven plasmas neutralized JR-CSF wild-type virus at an inhibitory dilution (ID50)>100, the minimal dilution probed. The distribution of ID50 ratios (mutant/wt) is shown for each mutant (center line: median; box limits extend from the 25th to 75th percentiles; whiskers indicate minimum and maximum values). A dotted line at ID50 ratio (mutant/wt)=1 indicates equal neutralization sensitivity of the mutant Env compared to the wildtype reference. Thresholds used for categorizing mutants into highly (ID50 ratio (mutant/wt)>10; colored red) and moderately (10>ID50 ratio (mutant/wt)>4; colored blue) sensitive phenotypes are indicated by dotted lines. **B.** IC50 values determined for each mutant against CD4 binding site (CD4-IgG2 and mAbs b6 and b12) and V3-crown (mAbs 1-79 and 447-52D) directed inhibitors were compared with median plasma ID50s as determined in A. Colorcode as in A. Titrations of each plasma and antibody on all viruses was done once, except for the JR-CSF wt reference for which plasma ID50s and nAb IC50s were derived by fitting a curve to pooled datapoints from n≥4 titrations.

Consistent with the limited cross-neutralizing activity observed in HIV-1 infection, JR-CSF wild-type (wt) was neutralized by only a subset of plasmas (11/18) (S1A and S1B Fig). While the majority of gp120 mutations did not lead to a relevant change in neutralization sensitivity, 23 mutants displayed a high (> 10-fold above wt) and 7 mutants a moderate (4-10 fold above wt) median increase against plasma (Fig 2A). Notably, these mutants also recorded high and modest neutralization sensitivity against plasma samples that were inactive against JR-CSF wt (S1A and S1B Figs) and across plasmas from individuals with different subtype of infection (S1C Fig), demonstrating a general release of conformational shielding in these 30 mutants. This was supported by the inhibition data with weak-nAbs. Only few mAb-mutant combinations deviated from this overall pattern of increased sensitivity, suggesting that in these cases the mutation directly affected the mAb epitope. Collectively, the 30 neutralization sensitive mutants recorded with a > 10-fold enhanced sensitivity to at least one V3-crown mAb (1-79, 447-52D) and one CD4bs directed inhibitor (b6, b12 or CD4IgG2) (Fig 2B and Table S2).

Known resistance inducing mutations were confirmed for some bnAbs. However, in contrast to weak-nAbs, bnAbs showed no or comparatively moderate increases in neutralizing potency across the mutant panel regardless of their specificity, with only a few bnAb-mutant combinations reaching a 10-fold increase, most of which fell within the 30 defined mutants with generally enhanced neutralization sensitivity (2G12: 2/2; PGT135: 1/1; 2F5: 9/10; 4E10: 9/9; 10E8: 3/5) (Fig 3A). Notably, the most consistent increase in neutralizing activity against these mutants was observed for MPER bnAbs known to exert their neutralizing activity primarily after CD4 binding when concomitant Env trimer opening increases MPER exposure [10, 27, 48]. Modest potency gains were also observed for 2G12 and for PGT135 which similar to MPER bnAbs, has been found to harbor post-attachment neutralization activity [10]. The CD4bs bnAb VRC01, the V2-apex bnAb PGT145 and several V3-glycan bnAbs (PGT121, PGT128, and PGT130) showed no increase in neutralization potency >10-fold across the entire panel.

**Fig 3.**
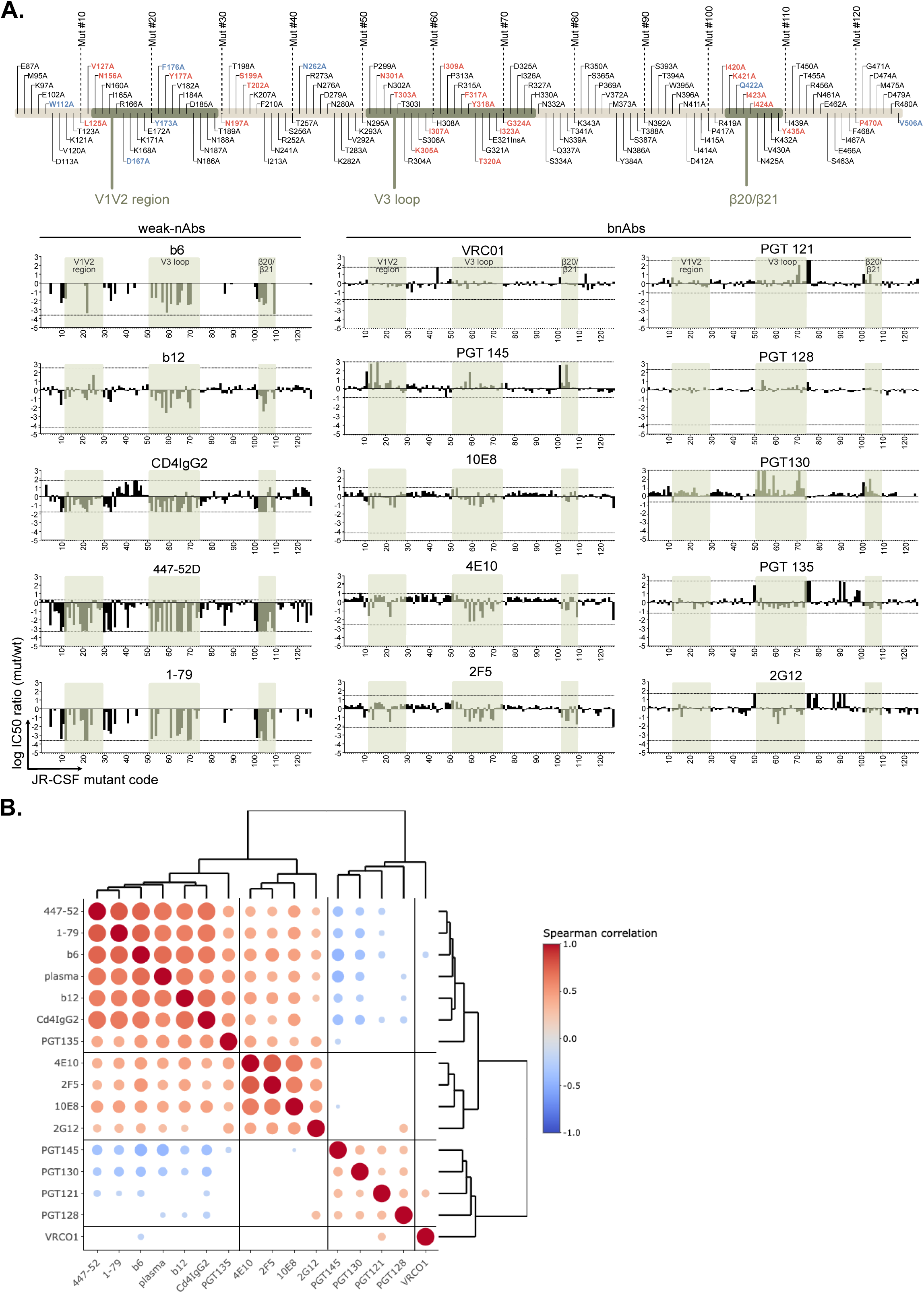
HIV-1 Env generalized neutralization sensitivity impacts differentially on the potency of broadly neutralizing antibodies (bnAbs) The whole JR-CSF Env mutant panel was tested for neutralization sensitivity against bnAbs targeting four major epitopes (CD4bs: VRC01; V3-glycan: PGT121, PGT128, PGT130, PGT135; 2G12; V2-glycan: PGT145; MPER: 2F5, 4E10, 10E8). A. Graphs show the log IC50 ratio (mutant/wt) for each Env mutant. IC50 ratios corresponding to minimal and maximal concentrations of inhibitors probed are indicated with lines. Weak-nAb data from Fig 2B is shown for comparison. The order of Env mutants and shaded regions of interest are indicated on top. Titrations of most antibodies on all viruses was done once (n=1), except for the JR-CSF wt reference (n≥4). mAbs PG128, PGT145, 10E8 and 2G12 were titrated twice (n=2) on all mutants. B. Spearman correlation matrix and hierarchical clustering of neutralization fingerprints based on IC50 data in A and Fig 2. For correlation with PWH plasma the median reciprocal ID50 ratio (mut/wt) over 11 JR-CSF wt neutralizing plasma (Fig 2A) was used. All significant correlations (p-value ≤0.05) are indicated with a circle with bigger circles indicating stronger support (lower p-values).

Distinct, general patterns of activity of weak-nAbs, bnAbs, and patient plasma across the mutant panel were revealed by Spearman correlation analysis (Fig 3B). Antibody reactivity patterns fell into four categories defined by hierarchical clustering: cluster I characterized by strong positive correlation amongst plasma, weak-nAbs and bnAb PGT135, cluster II comprising MPER bnAbs and 2G12 with low but consistent positive correlation to cluster I, iii) cluster III with V3-glycan bnAbs (PGT121, PGT128 and PGT130) as well as V2-glycan bnAb PGT145 showing negative correlations to cluster I and iv) bnAb VRC01 for which no consistent correlations with other clusters were detected.

The similarity of MPER bnAbs and PGT135 with plasma and weak-nAb inhibition patterns underscores that the activity of these Abs benefits from structural destabilization of the trimer. PGT135 and 2G12 target overlapping epitopes including glycans at positions N295, N332 and N386 [49, 50] suggesting that their simultaneous engagement by these two Abs is to some extent facilitated by enhanced conformational flexibility of the trimer. Conversely, a decreased activity of PGT145 against less stable trimers is expected as its quaternary V2-glycan epitope at the trimer apex depends on close vicinity of the gp120 protomers [51]. The data further suggested that PGT130, which like PGT121 and PGT128 binds the GDIR motif at the V3-base but differs in the dependency on glycans located in V1V2 (N137, N156) and V3 (N301, N332) [52, 53], may also benefit from a more restrained closed conformation.

Collectively, the reactivity pattern highlighted qualitative differences of bnAbs, and suggests that the most potent and broad bnAbs including VRC01, PGT121 and PGT128 show a comparatively high tolerance towards perturbation in Env conformational stability.

### Generalized increases in neutralization sensitivity link with de-stabilization of the closed prefusion Env conformation

To explore common antigenic features associated with a generalized neutralization sensitivity we probed five moderately and 22 highly neutralization sensitive Env mutants in a cell-based binding assay. Cell-expressed Envs were triggered by increasing doses of soluble CD4 (sCD4) and gradual opening of the trimer measured by the binding intensity of the prototypic CD4i mAb 17b that depends on trimer opening to access its epitope [17, 54–58] (Fig 4A). JR-CSF wt and the N332A mutant, which showed no increased neutralization sensitivity, were added for comparison. N332A and wt Env failed to bind 17b in absence of sCD4 trigger while the majority of the sensitive mutants (20/27) bound 17b (Fig 4B, S2A and S2B Fig). Neutralization sensitive mutants (7/27) that did not bind 17b in absence of sCD4 included mutants in residues 420-424 and Y435 which are known to impact on 17b binding to monomeric gp120 [56]. In none of these cases 17b binding was however completely abolished since all neutralization sensitive mutants displayed a notable increase in 17b binding upon CD4 triggering. Except for Y173A and P470A, which showed increased 17b binding at baseline, sCD4 dose escalation revealed that all other 17b neutralization-sensitive mutants (25/27) had an increased propensity to adopt an open conformation, consistent with a decreased stability of the closed pre-fusion conformation in these mutants (Fig 4B, S2A and S2B Fig).

**Fig 4.**
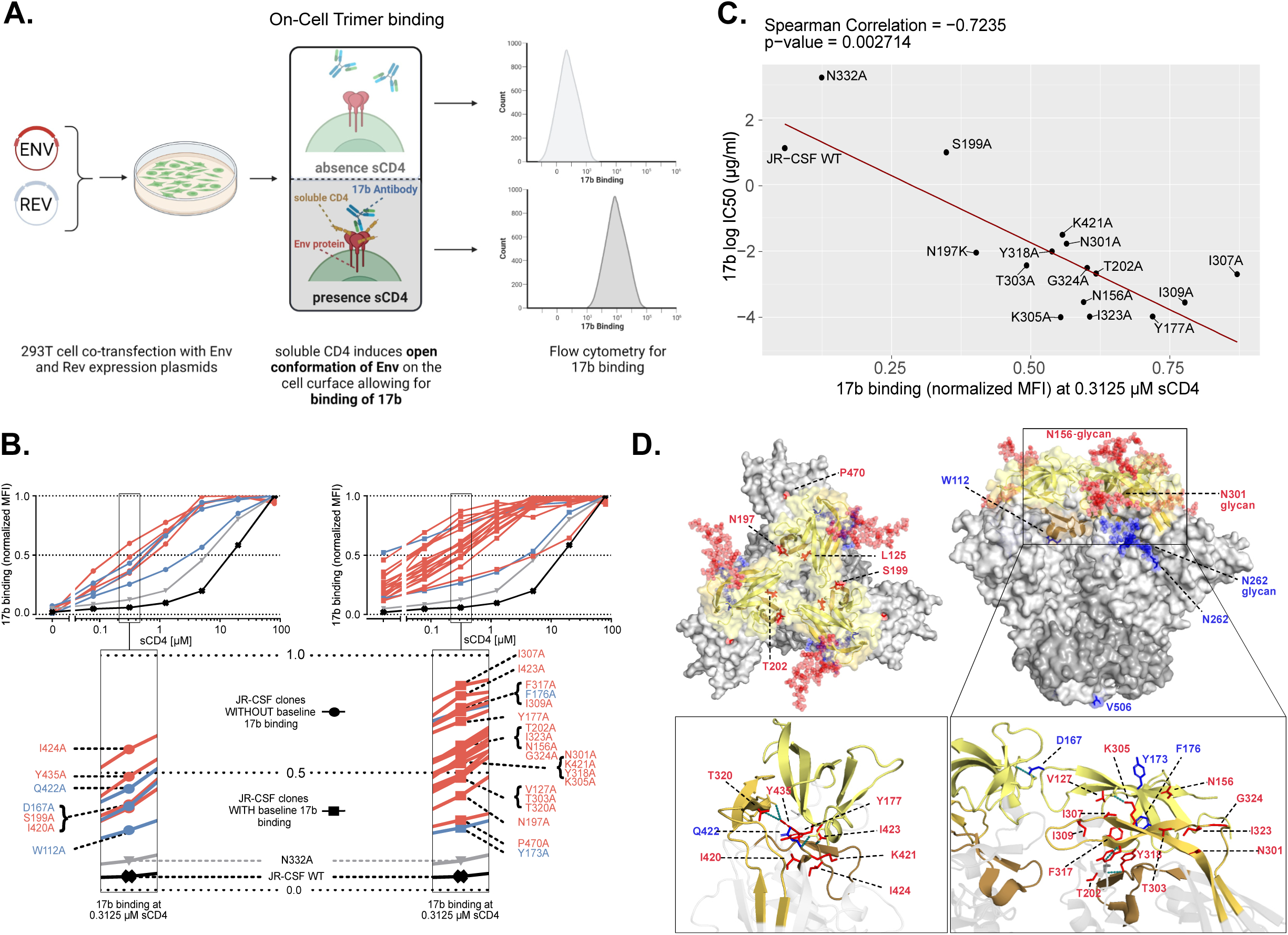
Characterizing the exposure of the co-receptor binding site by neutralization sensitive Env mutants. **A.** Schematic illustrating the experimental sCD4-induced opening of cell surface expressed Env leading to exposure of the co-receptor binding site. For detection by flow-cytometry mAb 17b was used as a co-receptor mimic. **B.** Opening of cell surface expressed Envs by increasing concentrations of sCD4 was monitored by staining with mAb 17b. Env mutants were devided into two groups showing enhanced 17b binding in absence of sCD4 (right) or not (left) compared to JR-CSF wt Env. The N332A mutant served as additional control. In the magnification on the bottom, the 17b staining is labelled for individual Env mutants. All titrations were performed once. Color code of mutant sensitivity as in Fig 2. **C.** Spearman correlation between 17b neutralization sensitivity of JR-CSF wildtype and mutant viruses and 17b binding according to B. Mutations directly affecting 17b binding [56] were not included (see also S2C Fig). **D.** Alanine-substitutions leading to moderate (blue) or high (red) generalized neutralization sensitivity of the JR-CSF Env were mapped onto the trimeric closed prefusion structure of the closely related JR-FL Env ectodomain (PDB: 5FYK; V1V2: yellow, V3: orange, β20-β21: brown, gp120: light grey, gp41: dark grey). Glycans on neutralization sensitive positions N156, N262 and N301 are depicted. The N197 glycan is not shown as JR-FL naturally lacks this PNGS. Enlargements on the bottom highlight several interactions (teal) sustained by residues in positions critical for neutralization sensitivity.

To link 17b exposure to neutralization sensitivity we measured the neutralization potency of 17b against 19 Env mutants with high general sensitivity alongside controls (S2C Fig).

Excluding mutants known to directly affect the 17b epitope (I420A, I423A, I424A, Y435A) [56, 59], the remaining mutants all showed high 17b sensitivity. 17b neutralizing activity and binding were tightly correlated confirming a propensity for trimer opening as a common trait of generalized neutralization sensitivity (Fig 4C and S2C Fig).

Mapping of the 30 JR-CSF Env mutations that confer generalized sensitivity on the crystal structure of the trimeric Env ectodomain from the closely related JR-FL strain (Fig 4D) highlights their positioning in domains relevant for conformational stability of the trimer [15, 16, 39]. The majority of mutations (26/30) were concentrated in and around V1V2, V3 and β20-β21 strands of the bridging sheet. The remaining four were mapped to C1 (W112A), C2 (N262A) and C5 (P470A and V506A). Glycans in gp120 contribute both to shielding and trimer stability [60]. JR-CSF wt contains 23 potential N-glycosylation sites (PNGS) within gp120. 17 of these PNGS were mutated in the full JR-CSF mutant panel (N=126; Fig 2, S1 Fig) and four of them (N156A, N197A, N262A and N301A) were associated with a generalized-sensitivity phenotype. As these residues are known to be important for Env stability [51, 61, 62], the gain in sensitivity upon loss of the glycosylation site may stem from both, a regained access to an epitope covered by this glycan or an increased propensity for trimer opening. Indeed, N197 has been shown to be critical for trimer integrity and neutralization resistance in several HIV-1 strains [28, 63–65] as the associated glycan is positioned to protect both V3 and the CD4bs [66, 67]. We also observed a dual role for the N301 PNGS in shielding and trimer stability. The decreased Env stability and increased neutralization sensitivity of N301A and T303A mutants was partially rescued in a T303I mutant, suggesting that the amino acid type in position 303 is also important (Fig 2, 3 and S2D Fig).

Since HIV-1 relies on the closed Env prefusion conformation to protect itself against antibody mediated neutralization [9, 11, 15, 16], amino acids regulating the maintenance of the closed state need to be highly conserved. To verify the conservation of the 30 generalized-sensitivity positions we identified for JR-CSF, we analyzed their conservation among group M Env sequences in the Los Alamos National Laboratory database (N=6084, Super filtered web alignment from year 2021 including Grp M and CRFs, http://www.lanl.com/). As expected we observed a high degree of conservation at these positions. 19 of the 30 Env mutations we investigated were recorded with >90% conservation (S3 Fig).

Focusing on mutations that confer high neutralization sensitivity in the context of JR-CSF, we next sought to confirm their sensitivity phenotype in a different Env genetic background. We chose for this the subtype A Env BG505 T332N known for its exceptional trimer stability [7, 38, 66, 68] and generated 20 variants carrying high sensitivity inducing mutations. Similar to JR-CSF, the majority of BG505 mutants (13/20) showed an enhanced propensity to undergo CD4-induced conformational transitions, exposing the V3-crown to a higher degree than BG505 T332N wt following sCD4 triggering (S4 Fig). However, in contrast to JR-CSF, neutralization sensitivity was increased only to CD4-IgG2 by these mutations. Other weak-nAbs and PWH plasma did not become more potent (S5 Fig). Thus, despite the high conservation of the positions impacting Env conformational stability, their mutation can entail different degrees of conformational destabilization and neutralization sensitivity depending on the virus strain.

### Neutralization fingerprinting using the JR-CSF Sensitivity Env mutant Panel (SENSE-19)

Considering the high conservation of the positions associated with generally increased neutralization sensitivity we reasoned that these JR-CSF mutants might be harnessed to characterize nAb activity. The use of these mutant Envs in neutralization screening may allow standardized typing of nAbs with respect to their preference and tolerance of Env structures. To ease incorporation of generally sensitive virus mutants into future nAb screens, we sought to focus on a concise collection of mutants with maximal information gain. Hierarchical clustering of all 126 JR-CSF mutants according to their neutralization sensitivity revealed a cluster of 18 mutants with high generalized sensitivity (S6 Fig). From this group mutant F317A was excluded due to relatively low infectivity. We additionally included mutants N197K and S199A as these target a PNGS with critical function in trimer stability and neutralization sensitivity ([28, 63–65] [66, 67]). We refer to this collection of 19 JR-CSF mutants as <u>Sens</u>itivity <u>E</u>nv Mutant Panel (SENSE-19).

We next used SENSE-19 to establish fingerprints of larger numbers of weak-nAbs and bnAbs building on those included in the full JR-CSF mutant screen (Figs 2 and 3). SENSE-19 fingerprinting of 11 additional Abs (5 CD4bs Abs, 3 V2-glycan Abs, 1 V3-glycan Ab, 1 gp120-gp41 interface Ab and 1 CD4i Ab) showed a pattern consistent with observations made on the full panel. Weak-nAbs displayed the expected strong increase in neutralization potency while bnAbs recorded comparatively small gains in activity or resistance (Fig 5A).

**Fig 5.**
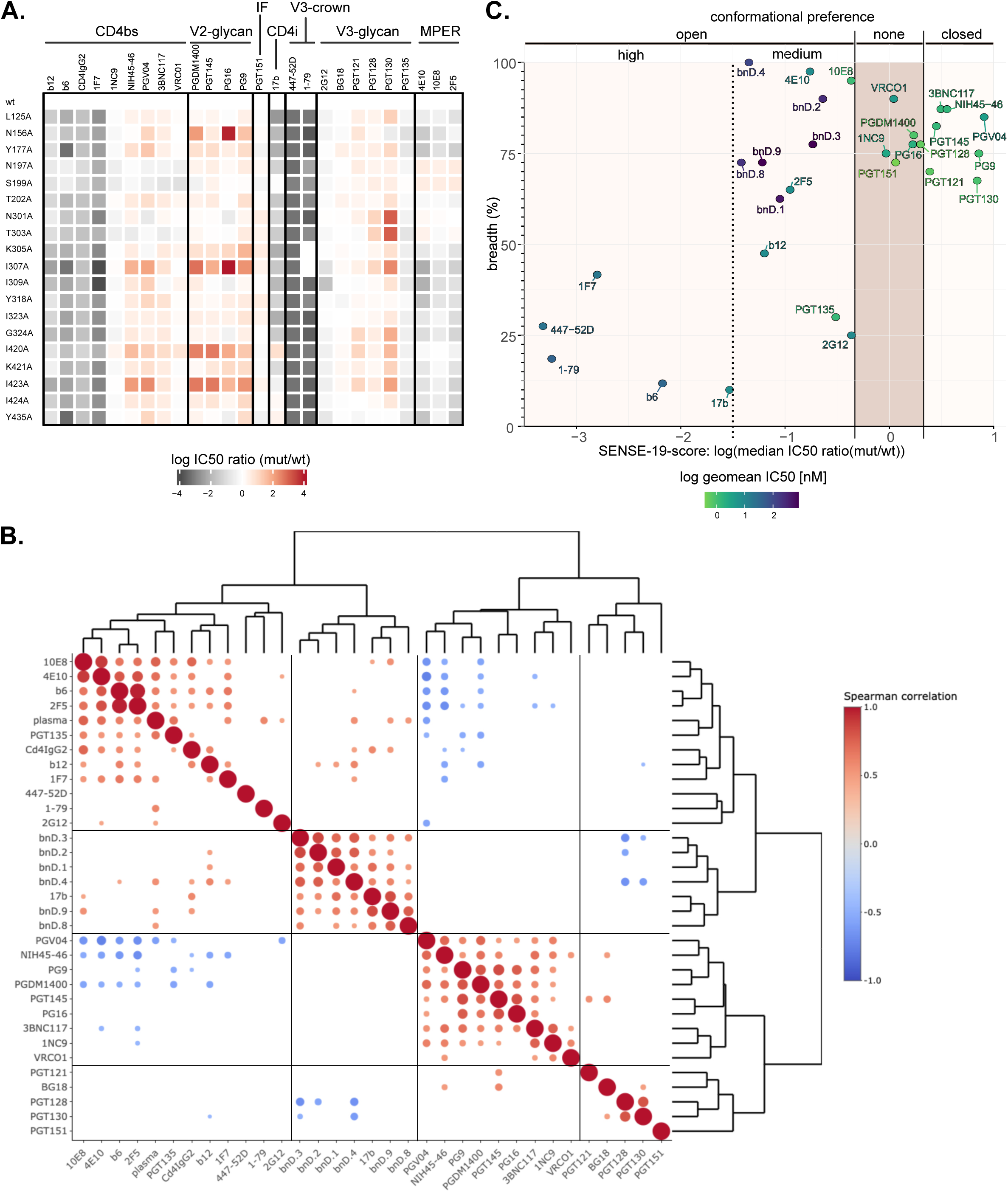
Neutralization fingerprinting of nAbs on a neutralization Sensitive Env mutant Panel (SENSE-19) Pseudovirus neutralization data on TZM-bl cells. **A.** IC50 ratios (mutant/wt) of an extended panel of nAbs were displayed as a heatmap. Abs are ordered by epitope. All titrations were performed once. **B.** Spearman correlation matrix and hierarchical clustering of SENSE-19 neutralization fingerprints from Abs, bnDs and PWH plasma (median reciprocal ID50 ratio (mut/wt) over 11 JR-CSF wt neutralizing plasma (Fig 2A)) All significant correlations (p-value ≤0.05) are indicated with a circle with bigger circles indicating stronger support (lower p-values). **C.** Comparison of three Ab quality parameters: neutralization breadth, potency and SENSE-19 score. Breadth and potency were determined on a cross-subtype 40 virus panel.

We further probed the performance of monovalent, broadly neutralizing DARPin (bnD) based inhibitors against SENSE-19. We included the V3-crown specific bnDs (bnD.1, bnD.2, bnD.3 and bnD.4; [44, 59]) and bnDs recognizing αV3C, an alpha-helix within the V3 C-terminal strand (bnD.8 and bnD.9; [59]). All six bnDs require Env opening for binding and exhibit dominant post-attachment neutralization activity [44, 59]. In agreement, the bnDs yielded notably enhanced potency across SENSE-19 (Fig 5C and S7 Fig). Interestingly, spearman correlation analysis and hierarchical clustering of SENSE-19 neutralization fingerprints revealed a cluster of conformation-specific inhibitors recognizing CD4-induced epitopes comprising Ab 17b as well as the bnDs tested (Fig 5B). This cluster was distinct from a second cluster comprising plasma and Abs with restricted activity against closed prefusion Env. While the latter cluster comprised a few bnAbs with dominant post-attachment activity, all other bnAbs clustered separately. Interestingly, SENSE-19 fingerprints of CD4bs and V1V2 bnAbs clustered together suggesting some similarity in conformational preference. V3-glycan bnAbs and the interface bnAb PGT151 formed a separate group (Fig 5B).

To differentiate nAbs and bnDs independent of potential direct interactions with residues mutated in the panel, we introduced a SENSE-19 score, defined as the median log IC50 ratio (mutant/wt) over the panel. Abs with a SENSE-19 score close to zero showed highest tolerance against changes in Env conformational stability. Particularly low variations in potency were evident for VRC01, 1NC9, BG18 and PGT151 demonstrating that Abs targeting different epitopes achieve high tolerance to structural changes. Classifying Abs and bnDs according to the SENSE-19 score together with neutralization potency and breadth, we observed four categories (Fig 5C). Abs with limited potency and breadth and strong dependence on trimer opening had SENSE-19 scores < −1.5. bnAbs and bnDs that benefit from trimer opening recorded a SENSE-19 score ranging from −1.5 to −0.3. bnAbs with high tolerance to structural changes displayed SENSE-19 scores between −0.3 to 0.3 and bnAbs depending on a closed trimer conformation scored >0.3.

Collectively, SENSE-19 fingerprinting provides insight of bnAb tolerance to structural changes and may be used in conjunction with a conventional neutralization breadth screen to define post-attachment acting bnAbs and broadly neutralizing inhibitors.

### Probing SENSE-19 mutants in the context of PBMC-based neutralization assays

An obstacle to the evaluation of neutralization activity remains the lack of standardizable, high-throughput assays to study HIV-1 inhibition in an environment similar to in vivo infection of primary CD4 T cells with replicating virus. Both the virus preparation and the variable infectivity of the target cells, peripheral blood mononuclear cells (PBMC) from random donors, are through-put limiting resources and can create a substantial assay-to-assay variability [69]. However, the use of PBMC-based neutralization in the evaluation of bnAbs has been reconsidered in recent years because the virus producer and target cells used may influence the neutralization efficacy recorded, at least for some Ab-virus combinations [69–76]. To verify whether SENSE-19 mutants record similar results in different assay systems, fingerprinting was performed in a PBMC based setup. We probed individual replication competent viruses in a conventional PBMC setup (Fig 6) and in a pooled virus library assay format similar to the approach used for Env mutational scanning (Fig 7) [77]. A requirement for both setups was to transfer SENSE-19 mutations into a full-length, replication-competent clone of JR-CSF (JR-CSF^rc^). In the conventional PBMC setup, virus infectivity in the supernatant was monitored. The library approach depends on sequence-based detection of the infecting strains. To identify individual strains by Illumina sequencing, we included 11 mutants from the SENSE-19 panel that fall within a stretch of DNA with a length that can be covered by Illumina sequencing (S10 Fig). As controls JR-CSF wt and four glycosylation site mutants (N295A, N332A, T450A and N461A) were included.

**Fig 6.**
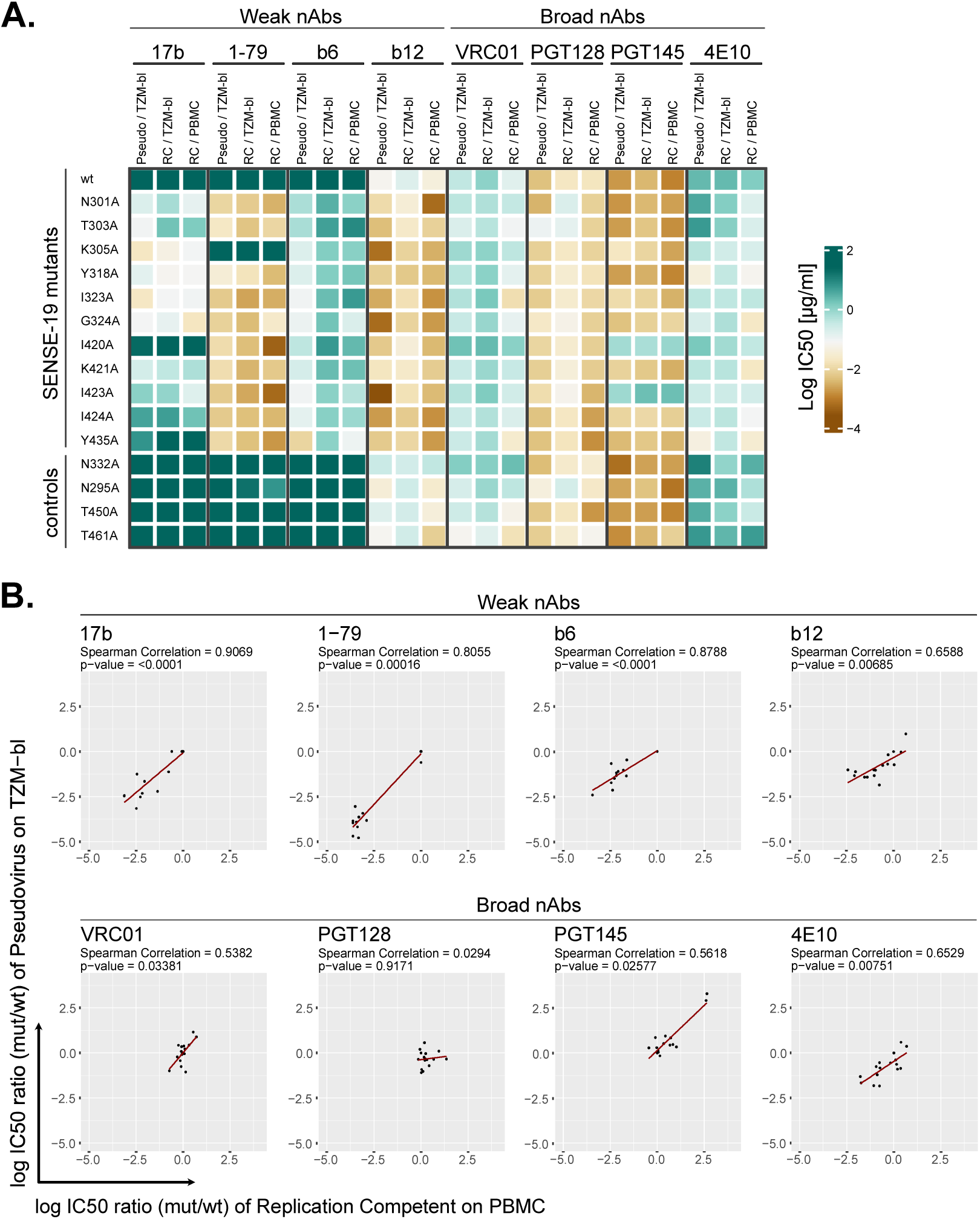
The generalized neutralization sensitivity phenotype is equivalent in different assay formats. JR-CSF wt virus and Env mutants were tested for neutralization sensitivity in three assay formats differing in the virus/cell combination: JR-CSF pseudovirus/TZM-bl, JR-CSF^rc^/TZM-bl, JR-CSF^rc^/PBMC. **A.** Heatmap comparing IC50 values from the three assay types. Titrations in the pseudovirus assay were performed once, except for the JR-CSF wt reference (n≥4). Titrations in the JR-CSF^rc^/TZM-bl assay were set up in triplicates. For the JR-CSF^rc^/PBMC assay results are shown from one of two independent experiments set-up in triplicate for each mAb respectively. **B.** Spearman correlation of shifts in neutralization sensitivity (log IC50 ratio (mutant/wt)) between the JR-CSF^rc^/PBMC and JR-CSF pseudovirus/TZM-bl assays (corresponding to data shown in S8A Fig).

**Fig 7.**
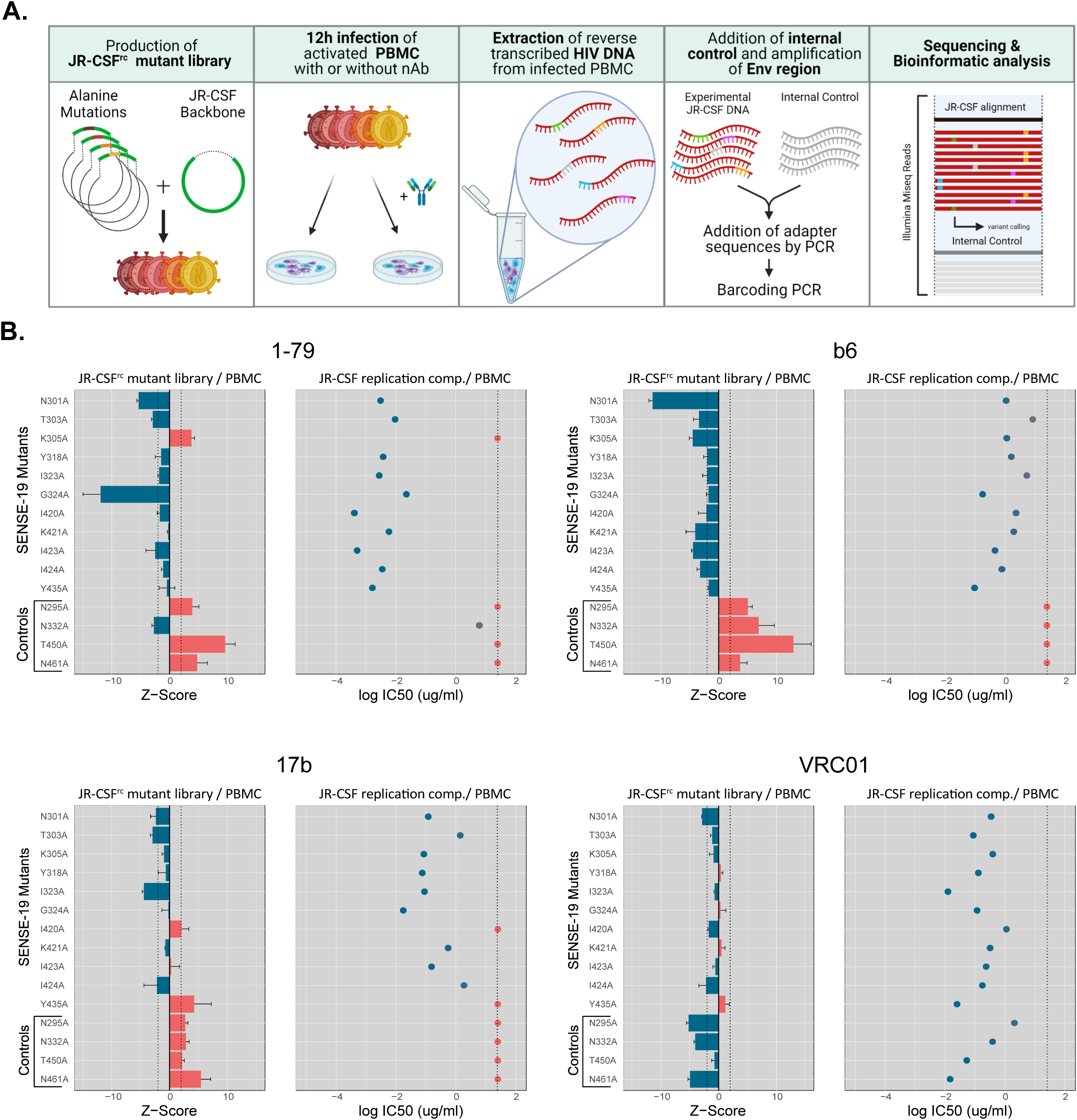
Validation of a virus mini-library neutralization assay for mAb fingerprinting with sequencing-based readout. The virus mini-library comprised 15 JR-CSF^rc^ mutant viruses including 11 of high generalized neutralization sensitivity and 4 PNGS mutants. Wt JR-CSF^rc^ was also included as reference. **A.** Schematic explaining the assay set-up and readout by illumina sequencing. **B.** Comparison of results from the virus mini-library neutralization assay with the PBMC based assay format testing JR-CSF^rc^ mutants individually for four different antibodies. Results indicating resistance and sensitivity to a nAb are colored red and blue respectively. Left panels for each Ab: Enrichment and depletion of mutant viruses in the mini-library in presence (25 µg/ml) versus absence of Ab are depicted as Z-scores (error bars: STD). A positive Z-score indicates enrichment and therefore resistance to the Ab, a negative Z-score indicates depletion and therefore sensitivity to the Ab (n=3). The dotted lines indicate the threshold at which the enrichment/depletion in presence versus absence of antibody exceeds two times the standard deviation of read frequencies in absence of antibody (Z-scores=2 and −2). Right panels for each Ab: Resistance of individually tested mutants in the PBMC-based assay was attributed if the IC50 was above 25 µg/ml (dotted line).

In a first step, JR-CSF^rc^ mutants were generated and probed in two conventional neutralization setups, on PBMC and on TZM-bl cells. The results were compared to the setup with JR-CSF mutants as pseudoviruses on TZM-bl cells (Fig 6). Overall, we observed a similar pattern for the three assay systems with much stronger variations in potency across the mutant panel for weak-nAbs compared to bnAbs (Fig 6, S8 and S9 Figs). Only modest differences between the PBMC based assay and the two TZM-bl assays were apparent. Notably, while IC50s for most Abs correlated between the three assays (S9 Fig), Abs tended to be less potent in the JR-CSF^rc^/TZM-bl setup (Fig 6A, S8B Fig). Moreover, IC50 ratios (mut/wt) correlated for most Abs between the JR-CSF/TZM-bl and JR-CSF^rc^/PBMC formats (Fig 6B). Correlations tended to be less strong for bnAbs (in particular PGT128) which likely is a result of the bnAbs low variability in potency paired with the inherent variability of PBMC-based assays.

To probe the JR-CSF mutants in the library approach (Fig 7) individually propagated virus stocks were pooled into a library and used to infect stimulated PBMC in presence and absence of nAbs. Unlike in conventional neutralization assays, where nAb activity is probed over a wide concentration range, a single, high-dosed concentration of mAb was used in this library neutralization assay (25 µg/ml). Twelve hours post infection, DNA was isolated from the PBMC cultures and the Env region chosen for monitoring was PCR amplified from reverse transcribed non-integrated viral DNA using barcoded primers. A DNA fragment derived from a plant-gene flanked by the same primer binding sites used for Env-enrichment was added to each sample as an internal control. The amplified DNA from antibody treated and non-treated samples together with the combined internal control was sequenced by Illumina Mi-seq, and individual reads were attributed to the internal control, different mutants or the wt Env using the previously established pipeline VirVarSeq [78]. The frequency of observed reads of each individual Env mutant were used to calculate Z-scores. A positive Z-score indicates relative enrichment of the Env in presence versus absence of a nAb and thus resistance, a negative Z-score indicates sensitivity (Fig 7). We probed the feasibility of this assay format with three weak-nAbs (b6, 1-79 and 17b) and bnAb VRC01. The library setup yielded results in agreement with the TZM-bl pseudovirus assays: Resistant mutants were enriched in the library in the presence of Abs 1-79, b6 and 17b while most SENSE-19 mutants were depleted or remained at similar level as when no Ab was present (Fig 7B). In contrast, all mutants were sensitive to VRC01 and by consequence none of them was enriched. We therefore conclude that the translation of the SENSE-19 screen to a library-based screen on PBMC is feasible and has potential for screens where a qualitative readout is useful. This approach may be particularly useful, when a large number of mutant and naturally evolved Envs are combined into a screening library. Overall, the comparison of PBMC and TZM-bl assay systems highlighted that screening with SENSE-19 mutants can be integrated in the assay format of choice and provides information on the tolerance of inhibitors to trimer opening.

## Discussion

In general, bnAbs are known for their capacity to engage the closed Env trimer but, in accordance with their activity against diverse viruses, must also harbor a considerable capacity to cope with the conformational plasticity of Env [20, 21, 79, 80]. Here, we systematically investigated the effect of Env conformational dynamics on bnAb activity. Generalized virus neutralization sensitivity has been associated with more frequent sampling of Env downstream conformations [17, 20]. In agreement, surveying the CATNAP database we found that weak-nAbs are more potent against Tier-1 viruses, which more frequently sample downstream conformations in the absence of CD4, compared to higher tiered viruses. Interestingly, some bnAbs with different epitope specificities also showed higher potency against Tier-1 viruses although this effect appeared less pronounced (Fig 1).

To capture the impact of Env conformational plasticity on Ab neutralization we sought to assemble Env mutants that can serve as indicator panel for neutralization screens. Screening 126 JR-CSF gp120 mutants we defined 30 mutants that displayed a generalized enhanced neutralization sensitivity against plasma from PWH and weak-nAbs, as seen with Tier-1 like Envs. These mutants displayed a lower stability of the closed conformation in comparison to wildtype. This lead to enhanced exposure of CD4-induced epitopes as demonstrated by cell-surface Env staining and neutralization sensitivity to Abs and inhibitors with post-attachment activity. While all neutralization sensitivity conferring mutations were located within gp120, they also enhanced sensitivity to bnAbs targeting the MPER which are known to require an open Env conformation for binding [8, 27, 30]. In agreement with the fact that prolonged periods of virus preincubation with MPER mAbs enhances neutralization [48], more frequent spontaneous sampling of open Env and/or altered entry process kinetics may lead to enhanced sensitivity to MPER bnAbs. More generally, spontaneous trimer opening and prolonged exposure of CD4-triggered conformations will benefit all inhibitors with post-attachment activity.

To provide a screening tool to monitor the effect of Env stability modulations on neutralization efficacy we introduced the SENSE-19 mutant panel which compares Ab activity against 19 JR-CSF Env variants with generalized neutralization sensitivity to wildtype JR-CSF (Fig 5). While bnAbs overall were less affected by increased trimer flexibility than weak-nAbs, scoring by the SENSE-19 panel provided novel insights on bnAb activity upon trimer opening. Increased potency against the SENSE-19 panel compared to JR-CSF wt was common among bnAbs with post-CD4 attachment activity. A second group of bnAbs tolerated SENSE-19 mutations well, with only modest potency changes compared to wt. Interestingly, this tolerance was achieved by bnAbs targeting different epitopes. A third group of bnAb reactivity was signified by a loss in activity on SENSE-19 in agreement with their dependence on a stable closed trimer conformation for optimal binding.

Collectively, the SENSE-19 screens provided compiled information on breadth, potency, and structural dependencies, and in particular, distinguished bnAbs with post-attachment activity. We postulate a utility of SENSE-19 as an extension to existing multi-subtype Env panels used for neutralization breadth and potency determination as it offers a framework for classifying bnAb activity based on structural changes in the HIV-1 Env protein. Virus quasispecies contain many variants with subtle sequence changes inflicting modest to large conformational changes and bnAbs used for prevention need to be able to combat them. We therefore consider the ability to cope with conformational plasticity as a feature of bnAbs that could provide relevant additional information about their breadth and thus provide an additional selection criterion to ascertain that the best bnAb or the best combination of bnAbs is moved forward in development.

## Materials and Methods

### Antibodies and Reagents

Reagents were kindly provided by following groups: Soluble CD4 (sCD4) comprising all four immunoglobulin domains [81, 82] and CD4IgG2 [83] by W. Olson (Progenics Pharmaceuticals Inc., Tarrytown, New York, USA); monoclonal antibodies (mAbs) VRC01, PGV04 [32] by J. Mascola (NIAID, NIH, Bethesda, USA); mAbs b12 [84], b6 [85], PG9 and PG16 [47], PGT121, PGT128, PGT130, PGT135 and PGT145 [31] and PGDM1400 [86] by D. Burton (The Scripps Research Institute, La Jolla, USA); mAbs 2G12 [87], 2F5 [88], 4E10 [89, 90] and 1F7 [91] by H. Katinger (Polymun, Vienna, Austria); mAbs 1-79, 3BNC117 and NIH45-46 [92] by M. Nussenzweig (Rockefeller University, New York, NY, USA); 447-52D [93] by S. Zolla-Pazner (Icahn School of Medicine at Mount Sinai, New York, NY, USA); 10E8 [30], 48D and 17b via the NIH AIDS Research and Reference Reagent Program, Division of AIDS, NIAID, NIH. Soluble CD4 containing the two amino-terminal immunoglobulin domains (sCD4-183) was expressed in E. coli and purified as described [25]. An overview is provided in Table S2.

### Human sera

All analyzed plasma samples (Table S1) were derived from specimens stored in the biobanks of the Swiss HIV Cohort study (SHCS) and the Zurich Primary HIV Infection Study (ZPHI). The Swiss HIV Cohort Study (SHCS) is a prospective, nationwide, longitudinal, non-interventional, observational, clinic-based cohort with semi-annual visits and blood collections, enrolling all HIV-infected adults living in Switzerland [94]. The SHCS is registered under the Swiss National Science longitudinal platform: http://www.snf.ch/en/funding/programmes/longitudinal-studies/Pages/default.aspx#Currently%20supported%20longitudinal%20studies. Detailed information on the study is openly available on http://www.shcs.ch.

The Zurich Primary HIV Infection study (ZPHI) is an ongoing, observational, non-randomized, single center cohort founded in 2002 that specifically enrolls patients with documented acute or recent primary HIV-1 infection (www.clinicaltrials.gov; ID NCT00537966) [95].

The SHCS and the ZPHI have been approved by the ethics committee of the participating institutions (Kantonale Ethikkommission Bern, Ethikkommission des Kantons St. Gallen, Comité départemental d’éthique des spécialités médicales et de médicine communautaire et de premier recours, Kantonale Ethikkommission Zürich, Repubblica e Cantone Ticino - Comitato Ethico Cantonale, Commission cantonale d’éthique de la recherche sur l’être humain, Ethikkommission beider Basel for the SHCS and Kantonale Ethikkommission Zürich for the ZPHI) and written informed consent had been obtained from all participants.

### Cells

293-T cells were obtained from the American Type Culture Collection. The TZM-bl cell line was obtained from NIH ARP. Both cell lines were cultivated in DMEM medium supplemented with 10% fetal calf serum (FCS) and 100 µg/ml streptomycin and 100 U/ml penicillin (P/S) (referred to as cell culture medium hereafter). All cell culture reagents were obtained from Gibco (Thermo Scientific, Waltham, USA).

PBMC were purified from buffy coats from anonymous blood donations from healthy individuals obtained by the Zurich Blood Transfusion Service (http://www.zhbsd.ch/) under a protocol approved by the local ethics committee. PBMC from three individual donors were CD8+ T-cell depleted using Rosette Sep cocktail (StemCell Technologies Inc.), pooled, split into three parts and incubated for three days with either 5 µg/ml PHA, 0.5 µg/ml PHA or anti-CD3 mAb OKT3 in RPMI 1640 medium (10% FCS, 10 U/ml IL-2, glutamine and 1% penicillin-streptomycin) [96].

### Viruses

The plasmid collection of JR-CSF env alanine mutants [31, 46, 47] and was a kind gift from D. Burton (The Scripps Research Institute, La Jolla, USA). The BG505.W6M.ENV.C2 [97] encoding source plasmid was obtained from the NIH AIDS Research and Reference Reagent Program. BG505 Env mutants and additional JR-CSF Env mutants were generated using the QuikChange II XL Site-Directed Mutagenesis Kit (Agilent, Santa Clara, USA) according to the manufacturer’s instructions. A BG505 Env mutant with V1V2 deletion was generated as described previously [11]. Pseudotyped virus production was performed by co-transfection of 293T cells with Env expression plasmid and HIV vector pNLluc-AM carrying a luciferase reporter cassette (provided by A. Marozsan and J. P. Moore) as described previously [74]. Information on the composition of a 40 virus panel used to assess mAb breadth and potency can be found in Table S3.

To generate replication competent mutant viruses, the JR-CSF Env encoding fragments containing the desired alanine substitutions were ligated in-frame into a replication competent JR-CSF infectious molecular clone (pYK-JRCSF obtained through the NIH ARP, catalog no. 2708[98]) using InFusion technology (Takara Bio Inc., USA) according to instructions by the manufacturer. Replication competent virus was produced by transfection of 293T cells with pYK-JRCSF based vectors analogous to the production of pseudotyped virus [74].

### Neutralization assay using Env-pseudotyped virus

The neutralization sensitivity of Env-pseudotyped viruses against mAbs and plasma from people living with HIV-1 (PWH) was evaluated on TZM-bl reporter cells in 384 well plates. Virus input was diluted with cell culture medium aiming for a luciferase reporter readout of 500’000-1’000’000 relative light units (RLU) per well measured on an EnVision luminometer (Perkin Elmer, Waltham, USA) in the absence of inhibitors. The antibody concentration (IC50) or reciprocal plasma titer (ID50) causing 50% reduction in viral infectivity were calculated by fitting the data to sigmoid dose–response curves (variable slope) using Prism (GraphPad Software). If 50% inhibition was not achieved at the highest or lowest concentration, a greater than or less than value was recorded.

Two different assay setups were established. To verify their comparability, PG128, PGT145, 10E8 and 2G12 were tested in both setups over the whole JR-CSF panel (N=126) and no systemic discrepancies were identified.

In the first set up, serial dilutions of inhibitors and PWH plasma in cell culture medium supplemented with 50mM HEPES were prepared robotically in 384 well plates (60 µl per well) (Corning, Corning, USA). This was followed by adding 20 µl per well of pseudovirus suspension. After 1-2 hours incubation at 37°C, 20 µl TZM-bl cell suspension at 150’000 cells per ml density in cell culture medium supplemented with 60 µg/ml DEAE-Dextran were pipetted robotically to each well and the plates were incubated for 48 hours at 37°C before proceeding to the readout of luciferase signal on an EnVision luminometer (Perkin Elmer, Waltham, USA) as described [74].

In an alternative assay format serial dilutions of inhibitors in cell culture medium (40 µl per well) were added to 40 µl of virus suspension in 384 well plates. After 1-2hours incubation at 37°C 50 µl of the virus-inhibitor mixture were added to pre-seeded TZM-bl cell cells (384 well plates, 6000 cells/well in 30µl culture medium supplemented with 26.6 µg/ml DEAE-Dextran) and the plates were incubated for 48 hours at 37°C before readout of the luciferase signal on an EnVision luminometer (Perkin Elmer, Waltham, USA) as described [74].

Neutralization data of bnD.1, bnD.2, bnD.3 and mAb 447-52D on the full JR-CSF mutant panel was reproduced from Friedrich et al [44]. Neutralization data of bnD.4, bnD.8, bnD.9 as well as mAbs PGT128 and 17b on strongly neutralization sensitive Env mutants was reproduced from Glögl et al [59].

### Neutralization assay using replication competent HIV

In addition to the pseudovirus neutralization assay performed on TZM-bl cells, replication competent virus was used to infect TZM-bl cells or PBMC in presence and absence of an inhibitor. To determine relative infectivity the individual replication competent virus stocks were first titrated on TZM-bl cells (seeded the previous day in a 384 well plate at 6000 cells/well density in 30 µl culture medium supplemented with 26.6 µg/ml DEAE-Dextran). The plate was incubated for 48 hours at 37°C before readout of the luciferase signal analogous to the procedure when using pseudovirus detailed above. As replication competent viruses are cytolytic for TZM-bl cells at high concentrations, this allowed to determine the dilution of the viral stock at which the TZM-bl cells survive for at least 3 days while ensuring high signal over background during readout. This virus dilution was not only used for inhibition assays of replication competent virus on TZM-bl cells, but also as input for the PBMC-based neutralization assay in 384 well plates.

For inhibition assays on TZM-bl cells, replication competent virus was titrated with inhibitor in triplicates, pre-incubated for 1h at 37°C and 30 µl/well of virus-inhibitor mix were then transferred onto TZM-bl cells seeded the previous day (384 well plate, 6000 cells/well in 30 µl culture medium supplemented with 26.6 µg/ml DEAE-Dextran). The plate was incubated for 48 hours at 37°C before readout of the luciferase signal analogous to the procedure when using pseudovirus detailed above.

For inhibition assays on PBMC, replication competent virus was titrated with inhibitor in triplicate, pre-incubated for 1h at 37°C and 30 µl/well of virus-inhibitor mix were then transferred onto PBMC (seeded at 1.5 x 10^6^ cells/ml density in 30 µl/well RPMI 1640 with glutamine supplemented with 10 % FCS, 100 U/ml IL-2, and 1 % penicillin-streptomycin). After incubation for 7 days at 37°C, plates were spun down at 450 g for 2 min and 30 µl/well of supernatant were transferred onto TZM-bl cells (seeded the previous day in a 384 well plate at 6000 cells/well density in 30 µl culture medium supplemented with 26.6 µg/ml DEAE-Dextran). After a further 24-72 hours of incubation infection was quantified by luciferase reporter readout as described for the pseudovirus assay to assess virus inhibition by antibody relative to wells without antibody. Additional wells which received the double amount of virus input in the PBMC-step were used as control to ensure that cytolysis was not affecting the assay.

### Sequencing based evaluation of neutralization assay with JR-CSF point mutant mini-library

To prepare the mini-library of replication competent JR-CSF Env point mutant viruses, individually prepared stocks of replication competent mutant viruses were mixed at equal volume. For the neutralization assay with this virus library PBMC were seeded at 3 x 10^6^ cells/ml in 250 µl RPMI 1640 with glutamine supplemented with 10 % FCS, 100 U/ml IL-2, and 1 % penicillin-streptomycin on 24-well culture plates. 125 µl of the virus library were preincubated either with or without 125 µl inhibitory agent diluted in cell culture medium aiming for a final concentration of 25 µg/ml after transfer of the mix onto PBMC. Each condition was set up in triplicates. The mixture of virus library and inhibitor was pre-incubated for 1h at 37°C and then transferred onto the seeded PBMC. 12 hours post-infection the cells were spun down at 16’000 g for 10 min and washed with PBS. The reverse-transcribed non-integrated viral cDNA was then extracted from PBMC using QIAprep Spin Miniprep Kit (Qiagen, Aarhus, Denmark). Samples were handled with frequent glove and pipette tip changing while any solution used was aliquoted to prevent cross contamination [99, 100]. As an internal control, 1 µl at 1 pg/ml of a custom synthesized 544 base-pair sized dsDNA fragment (Twist Bioscience, South San Francisco, California, USA) containing 500 base-pairs of a plant-gene (EMB2768 – Farinopsis Salesoviana) flanked on each side by one of the two primer binding sites used for amplification of Env-fragments (described below) was added to each sample. In a first PCR (see table S4) amplification of Env fragments covering all alanine substitutions in the virus mini-library (and the internal DNA control fragment) was done with Env specific forward and reverse primers of three different lengths containing adapter sequences at the 5’ end.

Forward primers: CTTTCCCTACACGACGCTCTTCCGATCT(N)_6,7,8_AACCATAATAGTACAGCTGAATGAATC

Reverse primers: GGAGTTCAGACGTGTGCTCTTCCGATCT(N)_6,7,8_TCTGAAGATCTCGATCTCACTCT

The differing length of the primers artificially increases diversity of the otherwise highly similar amplicons for efficient MiSeq sequencing. In addition, 5 random nucleotides between the adapter-region and binding region of the primers were added to facilitate cluster generation during illumina amplification of reads. The forward and reverse primers of different lengths were used at 3.33 µM concentration each, to result in 10 µM forward and reverse primer mixtures.

The DNA was then purified from the PCR reactions using AMPure XP beads (Backman Coulter Life Sciences, Indianapolis, USA) at a 1:1 ratio (vol/vol) as recommended in the instructions by the manufacturer. After the clean-up, a 2^nd^ PCR was performed to barcode the individual samples using Index primers (TruSeq HT Kits, Illumina Inc., San Diego, California, USA) able to bind the adapter-region established during the previous PCR.

After PCR dsDNA concentration in samples was measured with the QuantiFluor system (Promega). The samples were subsequently diluted to 4 nM concentration and twenty-four samples were mixed together in equal parts. The Ilumina sequencing used a MiSeq Reagent Kits v2 (500 cycles) and was executed based on manufacturer’s instructions.

For evaluation individual reads were attributed to the internal control, different Env mutants or the wt Env using VirVarSeq [78]. Then, the relative frequencies of each Env variant (rF(variant y) _rep z_) computed by VirVarSeq (controlled for position-specific read quality) in samples treated or not with antibody were transformed into adjusted relative frequencies (arF (variant y)_rep z_) taking into account the number of reads corresponding to the internal control.

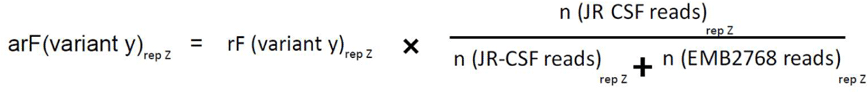

rep z: replicate 1, 2 or 3

n (JR-CSF reads): total number of reads aligned to JR-CSF reference by VirVarSeq

n (EMB2768 reads): total number of reads aligned to plant reference by VirVarSeq

Based on the adjusted relative frequencies of reads, Z-scores were calculated for individual Env variants of each experimental replicate as follows:

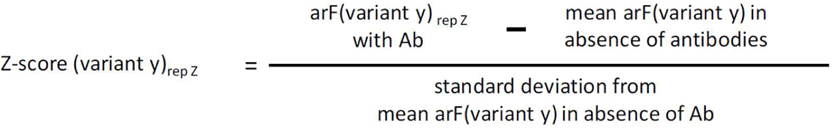

Wherein the mean adjusted relative frequency of an Env variant in absence of antibody is:

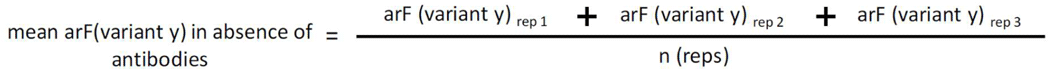

The internal control allows the comparison of read frequencies from Env variants in presence and absence of Ab and is particularly important in case the majority of viruses in the mini-library is sensitive to the Ab.

### Flow cytometric analysis of antibody binding to cell surface-expressed HIV-1 Env

10^5^ 293T cells per well were seeded in 1 ml cell culture medium in 12-well tissue culture plates and incubated at 37 °C. 24 hours later, cells in each well were transfected with a total of 1 μg DNA (Env-expression plasmid and pCMV-*rev* expression helper plasmid in 4:1 ratio) mixed with 3 μg 25 kDa linear PEI (Polysciences Inc., Warrington, USA) in 200 μl 150 mM NaCl. After settling the DNA-PEI complexes by a short spin (3 min., RT, 300 g) the cells were incubated 36 hours at 37°C. All subsequent steps were carried out at room temperature. Cells were harvested, wells transfected with the same Env-plasmid pooled, and distributed into 96-well round bottom tissue culture plates as individual staining reactions (cells from one 12-well were split into four staining reactions). For each staining reaction cells were washed once with 200 μl FACS buffer (DPBS (Gibco) with 2% FBS, and 2 mM EDTA) and stained for 20 min in 15 μl of FACS buffer with 10 μg/ml of primary antibody in the presence or absence of sCD4-183. After washing twice with 200 μl FACS buffer the cells were incubated in 30 μl of FACS buffer for 20 min with 1:1000 diluted APC-conjugated F(ab’)₂ fragment goat anti-human IgG (Jackson ImmunoResearch, West Grove, USA). Following two further washes with FACS buffer the cells were resuspended in 100 μl FACS buffer with 0.1 µg/ml propidium iodide (BD Pharmingen). Flow cytometry analysis was performed on a FACSVerse system (BD biosciences, San Jose, USA). Arithmetic mean of APC fluorescence intensity was calculated for Propidium Iodide negative single cell population as a measure of primary antibody binding to the live cells in each staining reaction.

Env mutants were devided into two groups showing enhanced 17b binding in absence of sCD4 or not compared to JR-CSF wt Env (threshold 10% of normalized maximal staining in presence of sCD4).

### Bioinformatics and Programs

To analyze Ilumina sequencing data from the JR-CSF library, VirVarSeq [78] was used to calculate the frequency of wildtype and mutant Envs as well as the internal control in presence or absence of an inhibitor. Sequences were trimmed using SeqTK [101] to remove residual, random nucleotides introduced to support Miseq amplification of reads.

The figures 1, 2, 3C,4C, 5, 6, 7B and supplemental figures S1, S2C, S3, S5, S7, S8 were produced using the R packages ggplot2 [102], heatmaply [103], tidyverse [104] and Hmisc [105]. Figures 3A, 4B and supplemental figures S2D and S4B were produced in GraphPad Prism 10 (GraphPad Software, Boston, Massachusetts, USA). FACS-generated data was processed using FlowJo^tm^ Software v10.9 (BD Life Sciences). Figure 4D was designed using PyMOL v2.5 (https://pymol.org/2/). Biorender (Subscription model, Canada) was utilized to make figures 4A and 7A. Affinity Designer (SerifEurope, UK) was used to finalize all figures.

## Data Availability Statement

All relevant data are within the manuscript and its Supporting Information files.

## Acknowledgements

The SHCS data are collected by the five Swiss University Hospitals, two Cantonal Hospitals, 15 affiliated hospitals and 36 private physicians (listed in http://www.shcs.ch/180-health-care-providers). We thank the patients participating in the ZPHI and the SHCS and their physicians and study nurses for patient care. We thank Therese Uhr for technical assistance and Zhaozhi Sun for his contributions in the frame of this master thesis. We are grateful to Merle Schanz and Magdalena Schwarzmüller for assistance with figure layout.

## Author contributions

AT directed this work. AT, PR and NF designed the study. CF, NF, BI, ES, UK and JW performed experiments. CF, NF, BI, ES, CM, DS, CP and PR analyzed data. HFG managed the SHCS and ZPHI cohorts, contributed patient samples and analyzed patient related data. CF, NF and AT wrote the manuscript, which all co-authors commented on. CF and NF contributed equally to the study. BI and ES contributed equally to the study.

## Supporting information captions

**S1 Fig. Characterizing the general neutralization sensitivity of JR-CSF envelope point mutants** The general neutralization sensitivity of 126 JR-CSF envelope (Env) mutant pseudoviruses was determined in a TZM-bl based assay using **A.** a set of 16 PWH plasma from chronic infection (eleven subtype B infected, five with non-B subtypes). **B.** ID50s for a subset of 5/16 plasma that did not inhibit JR-CSF wt Env at an inhibitory dilution ID50=100, the minimal dilution probed (four plasma samples from subtype B, one from non-subtype B infected individuals). **C.** ID50s for five plasma samples from PWH chronically infected with non-subtype B HIV-1 (subtype of infection in parentheses). Three plasma samples neutralized JR-CSF wt, two did not. (Boxplots show center line: median; box limits extend from the 25th to 75th percentiles; whiskers indicate minimum and maximum values). Mutants with high and moderate general neutralization sensitivity as assigned by the analysis presented in Fig 2A are colored in red and blue respectively. Table S1 provides an overview of the plasma samples used. Titrations of each plasma on all viruses was done once, except for the JR-CSF wt reference (n=4).

**S2 Fig. Characterizing the exposure of the co-receptor binding site by neutralization sensitive JR-CSF Env mutants** Opening of 293T cell surface expressed Envs by increasing concentrations of sCD4-183 is monitored by flow cytometry after staining with mAb 17b, directed against a CD4-induced epitope similar to the CCR5 co-receptor binding site. **A.** Example to illustrate the gating strategy applied during analysis by flow cytometry. **B.** Histograms showing the 17b staining obtained for each Env mutant. **C.** Spearman correlation between 17b neutralization sensitivity of JR-CSF wildtype and mutant viruses and 17b binding according to Fig 4B. In comparison to Fig 4C mutations directly affecting 17b binding [56] were included. **D.** Env mutants T303A and T303I are differentiated by 17b binding at baseline, i.e. in absence of sCD4, and by their propensity to expose the 17b epitope with increased concentrations of sCD4-183. All titrations were performed once.

**S3 Fig. Conservation of all 125 amino-acid positions mutated in the full JR-CSF Env mutant panel.** A superfiltered web alignment (available from the Los Alamos National Laboratory database, http://www.lanl.com) of diverse HIV-1 Env sequences (6084 Env sequences from group M including CRFs) was downloaded for the most recent year available (2021) and all relevant amino acid positions were analyzed for their conservation Residues associated with high (red) and moderate (blue) general neutralization sensitivity are indicated.

**S4 Fig. Analyzing the propensity of BG505_T332N (wt) envelope and mutants to adopt the CD4-induced conformation.** Titration of 293T cell surface expressed BG505 wt envelope and mutants with sCD4-183. The induced opening of the Env trimer was monitored with V3-crown directed mAb 1-79 and analyzed by FACS. **A.** Histograms showing mAb 1-79 staining. B. Dose-response curves for all Envs tested. The reference envelope BG505_T332N and the R308A mutant (the corresponding JR-CSF mutant has a wt-like phenotype) are colored in black and grey, respectively. BG505 mutants in red reproduce the enhanced propensity to adopt the CD4i state observed for corresponding JR-CSF mutants, the mutants in purple do not. A V1V2 deleted BG505 T332N envelope was included as control (dashed red line). K305A and I307A are 1-79 knock-out mutants (brown). MuLV envelope was used as control for unspecific staining (brown). The two panels show the same data without (left) and with (right) normalization to the highest signal obtained for each mutant respectively.

**S5 Fig. Probing the neutralization sensitivity of selected BG505 envelope mutants.** Neutralization sensitivity was determined for 17 alanine mutants based on the BG505 T332N Env (designated as wt) corresponding to JR-CSF alanine mutants with strong general neutralization sensitive phenotype using a pseudovirus assay system with TZM-bl cells. **A.** and **B.** Plots show changes in IC50 values relative to the wt envelope. A BG505 T332N Env lacking V1V2 (ΔV1V2) was included as a sensitive control. Depicted are results using **A.** CD4bs-directed, **B.** V3-crown directed (1-79), and V2-glycan directed (PG16, PGT145) Abs. **C.** ID50 values were determined for three BG505 Env mutants using 16 PWH plasma samples (13 infected with subtype B (circles) and 3 with non-subtype B (triangles) HIV). The minimal dilution of plasma tested was 1/100. For each mutant the median ID50 is indicated with a red bar.

**S6 Fig. Hierarchical clustering analysis of JR-CSF mutant Envs** All 126 JR-CSF mutant Envs included in the full virus panel were grouped according to their changes in neutralization sensitivity compared to wt (log IC50 ratios (mut/wt)) against the antibodies indicated at the bottom. For PWH plasma the median reciprocal ID50 ratio (mut/wt) over 11 JR-CSF wt neutralizing plasma (Fig 2A) was used.

**S7 Fig. Neutralization fingerprinting using a panel of generally neutralization sensitive Env mutants (SENSE-19)** Neutralization fingerprinting of monovalent bnDs and bivalent bnD-Fc constructs. IC50 ratios (mutant/wt) are displayed as a heatmap.

**S8 Fig. Probing the equivalency of the generalized neutralization sensitive phenotype in different assays formats** JR-CSF wt virus and Env mutants with and without high overall neutralization sensitivity were tested in three different assay formats. IC50 values were determined in a pseudovirus assay system with TZM-bl reporter cells and in two set-ups with replication competent virus (JR-CSF^rc^) infecting either TZM-bl cells or PBMCs. **A.** Heatmap of log IC50 ratio (mut/wt) values. **B.** Distribution of IC50 values corresponding to data in A. Box plot indicates median (center line), 25^th^ to 75^th^ percentiles (box limits) as well as minima and maxima (whiskers). Titrations in the pseudovirus assay were performed once, except for the JR-CSF wt reference (n≥4). Ab titrations in the JR-CSF^rc^/TZM-bl assay were set up in triplicates. For replication competent virus on PBMCs results are shown from one of two independent experiments set-up in triplicate for each mAb respectively.

**S9 Fig. Spearman correlations of virus neutralization sensitivity data from three different assay formats** Log IC50 values were correlated for the Abs indicated from two of three different assay formats respectively, based on data shown in Fig 6A.

**S10 Fig. Sequencing-based evaluation of an inhibition assay using a library of replication competent viruses on PBMC** The schematic indicates the positioning of primer binding sites used to determine the composition of a replication competent virus mixture containing JR-CSF wt and point mutated envs by Illumina MiSeq sequencing.

**Table S1** Overview on the characteristics of plasma samples from people living with HIV (PWH) used in the study

**Table S2** Overview of origin and source for all mAbs used in the present study

**Table S3** Information on the composition of a 40 virus panel used to assess mAb breadth and potency

**Table S4** PCR conditions used during sequencing-based evaluation of virus inhibition assays

