## Supplementary figures and images for "Assessing bnAb potency in the context of HIV-1 Envelope conformational plasticity"

### Supplemental figures S1-S10

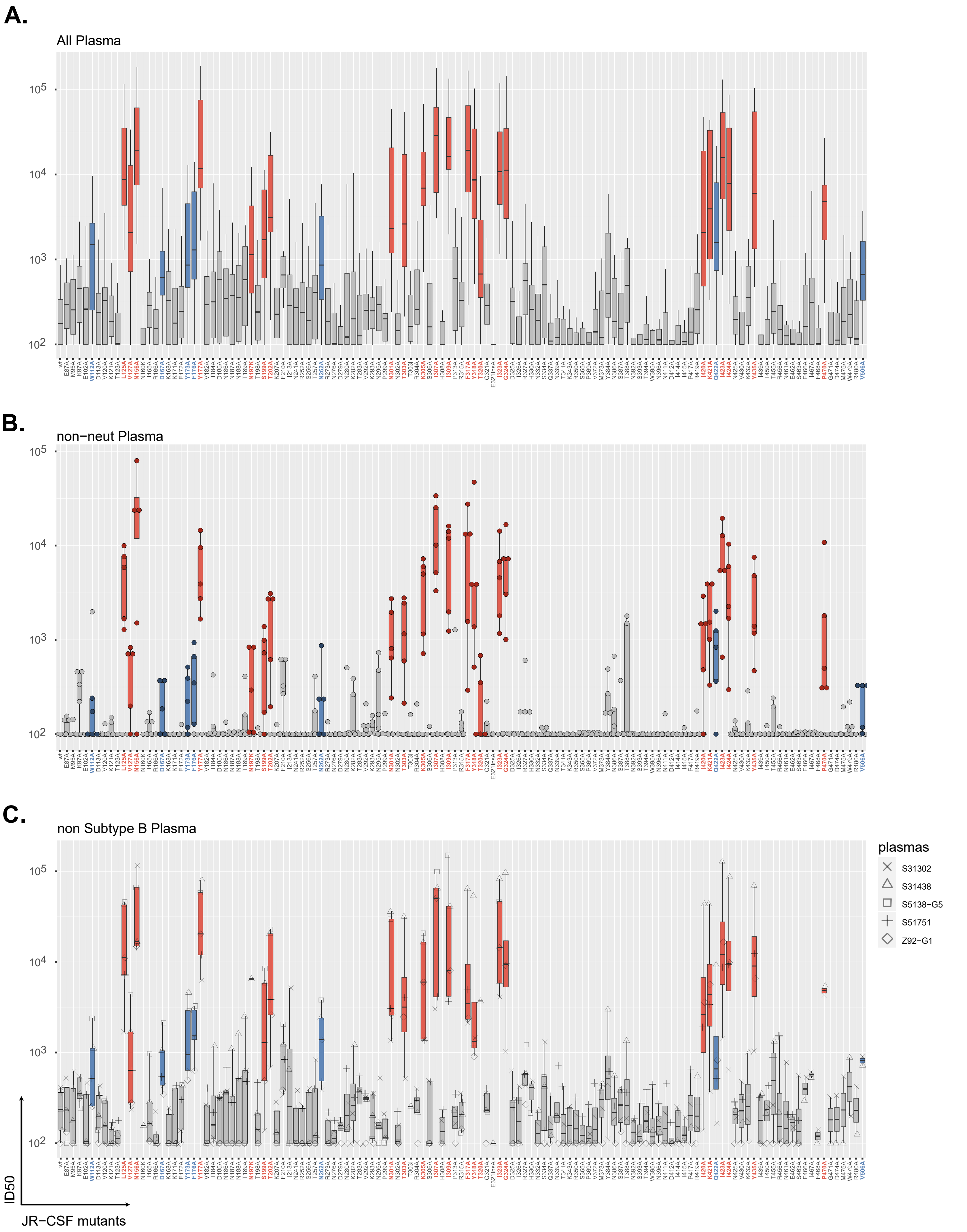

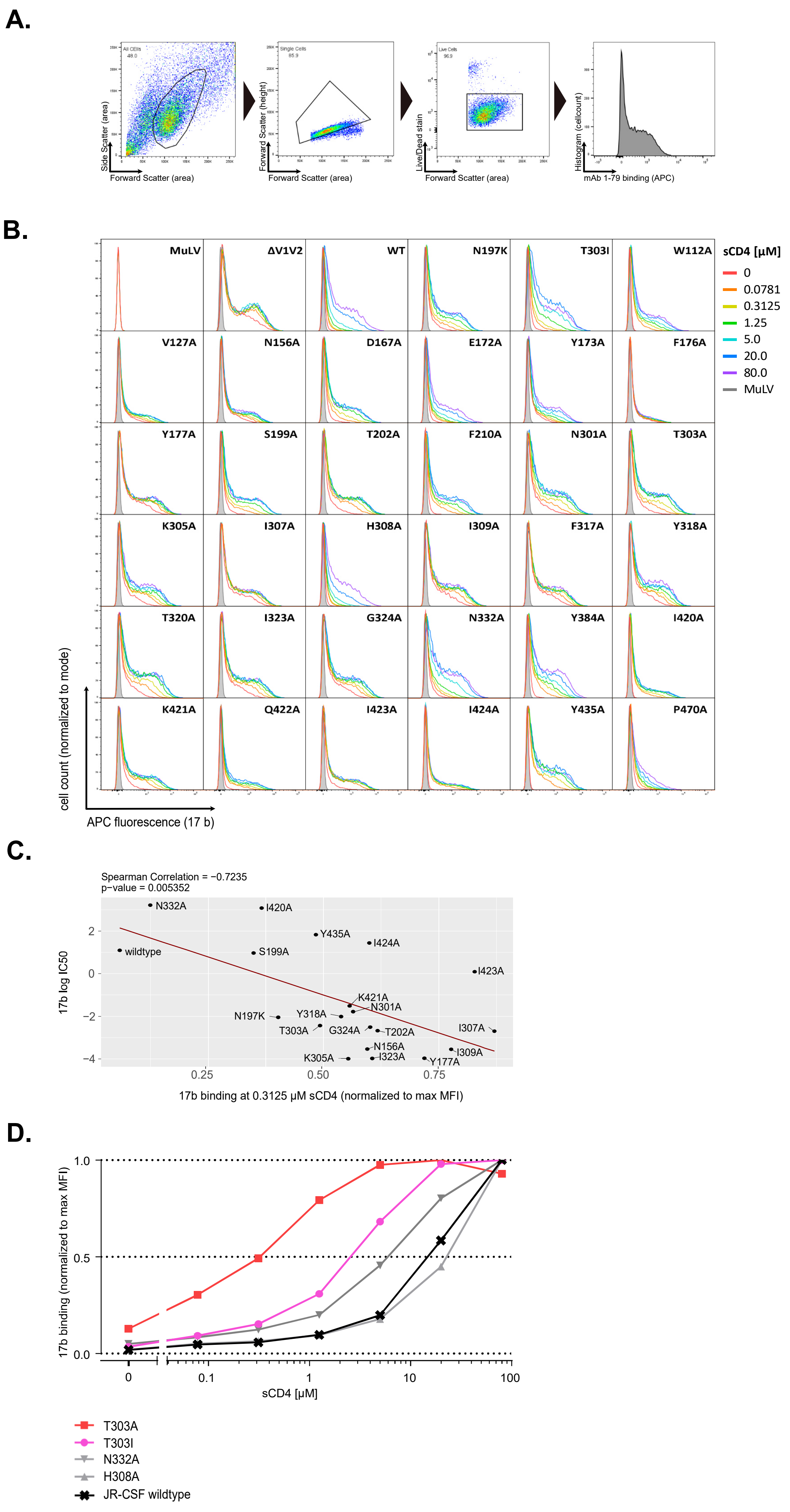

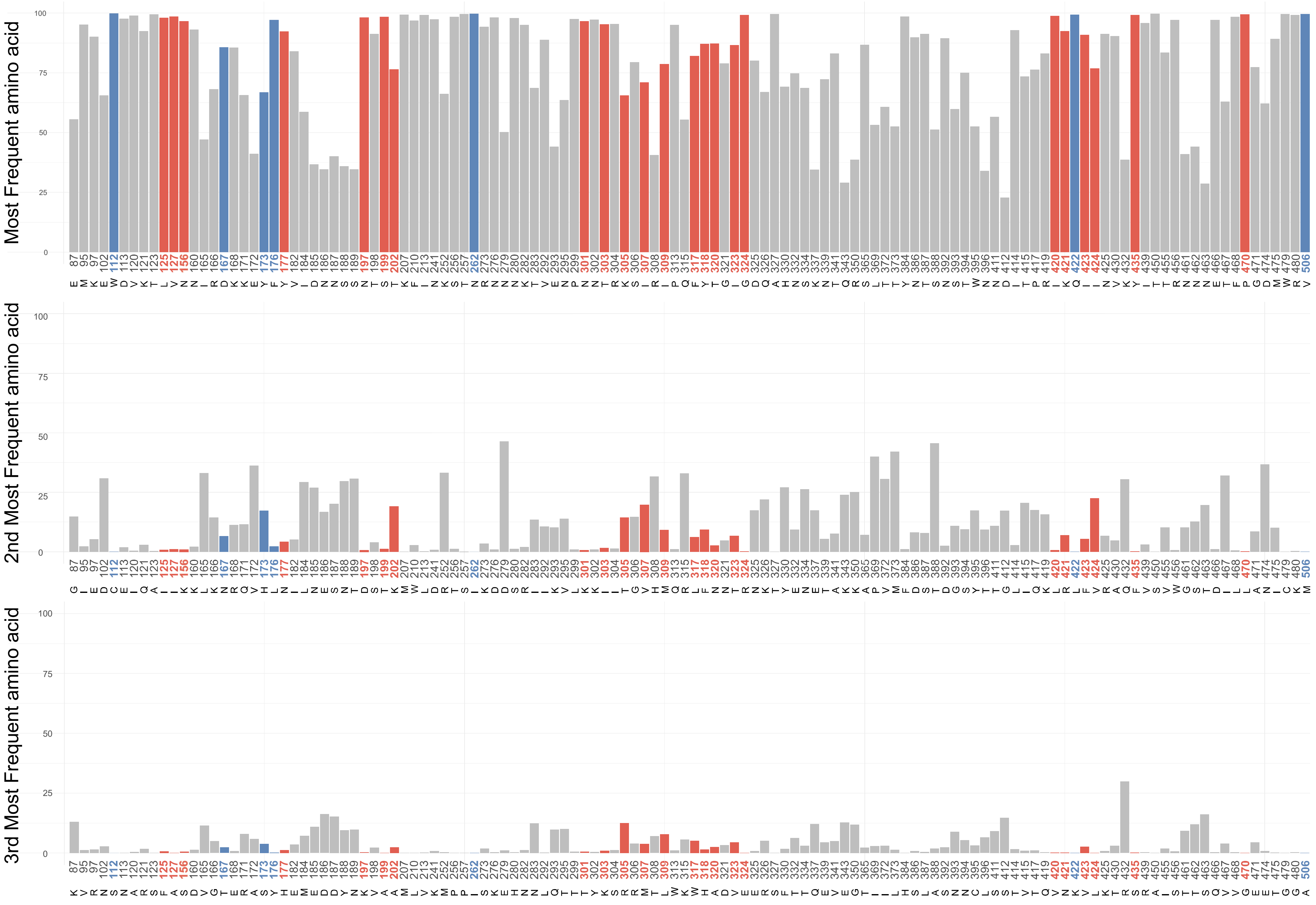

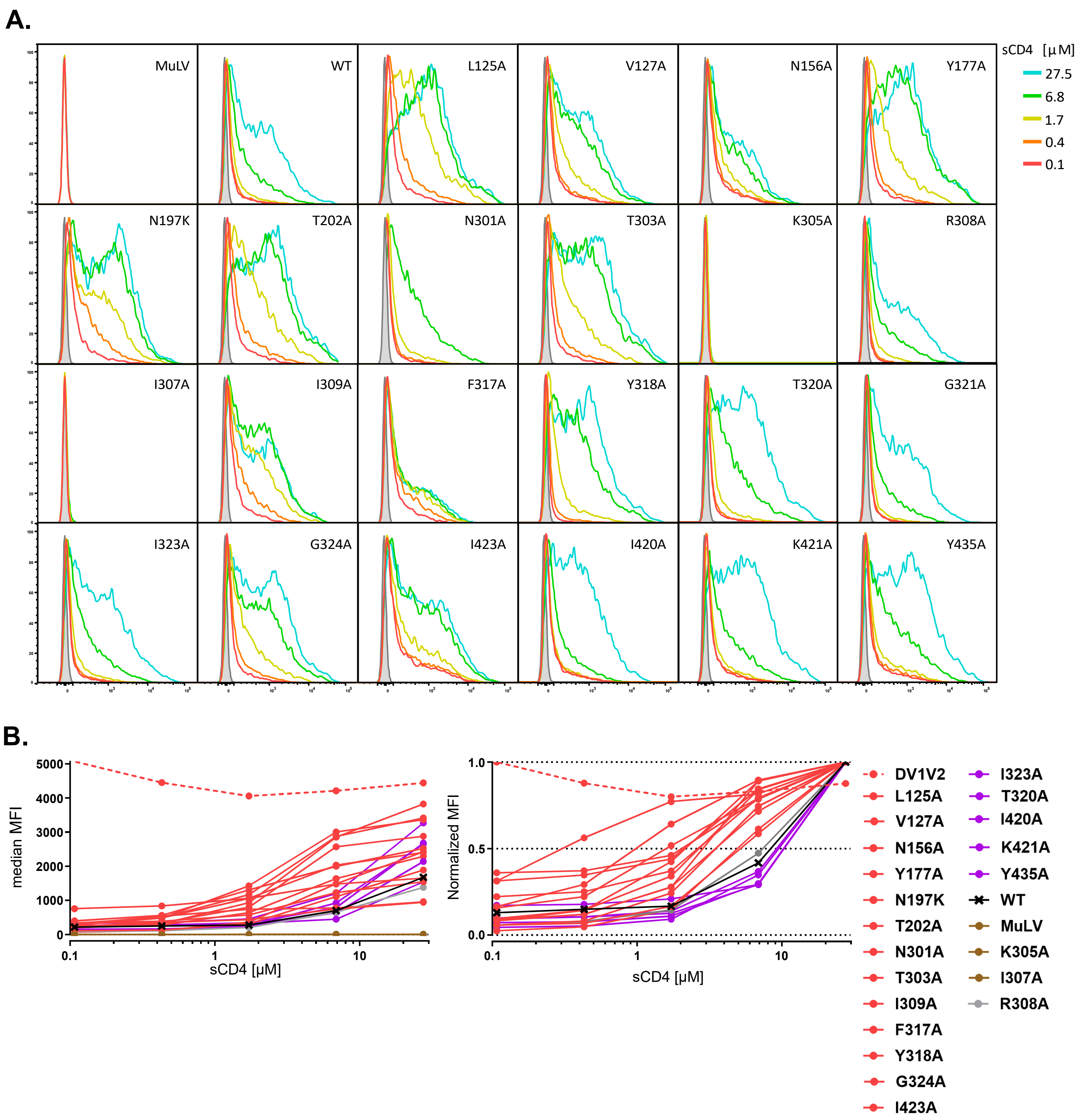

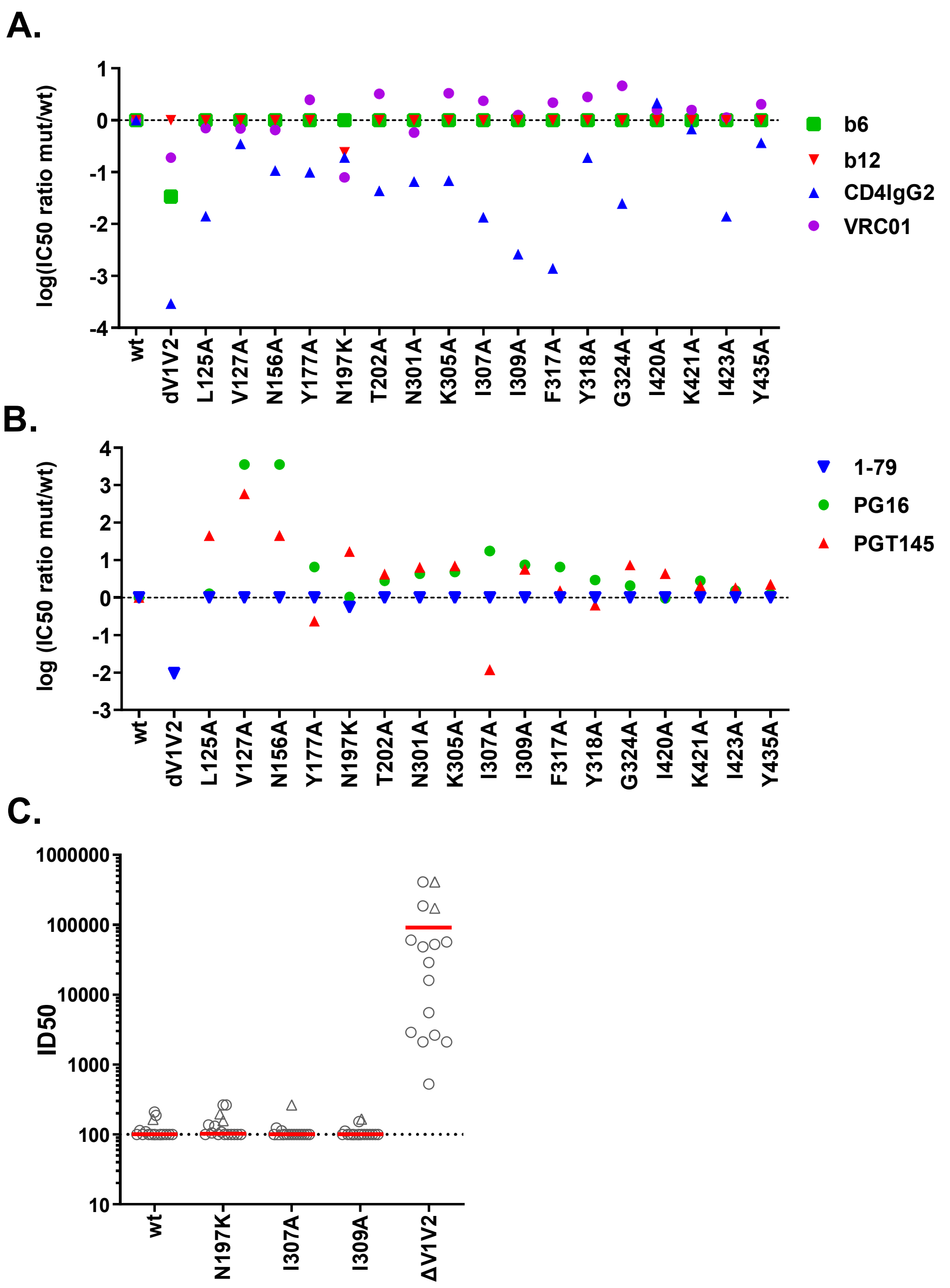

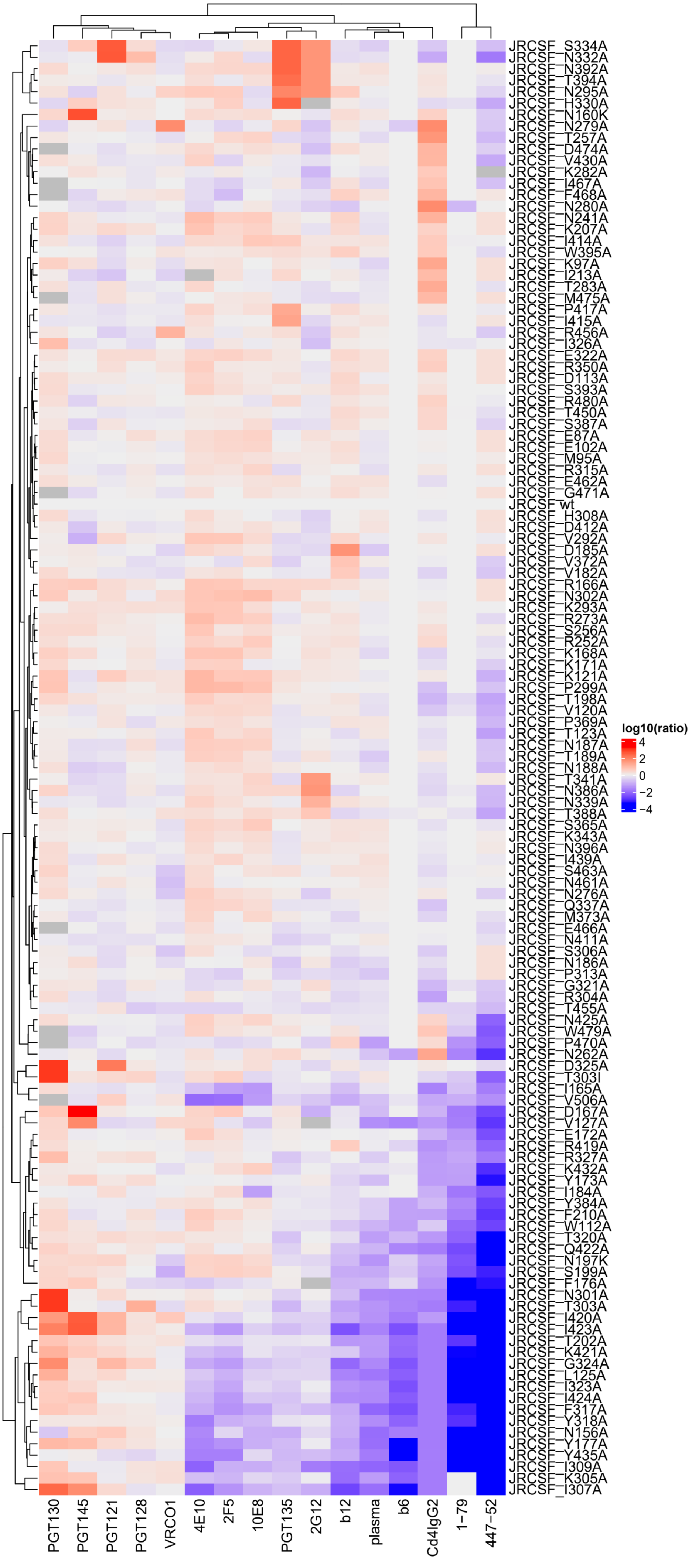

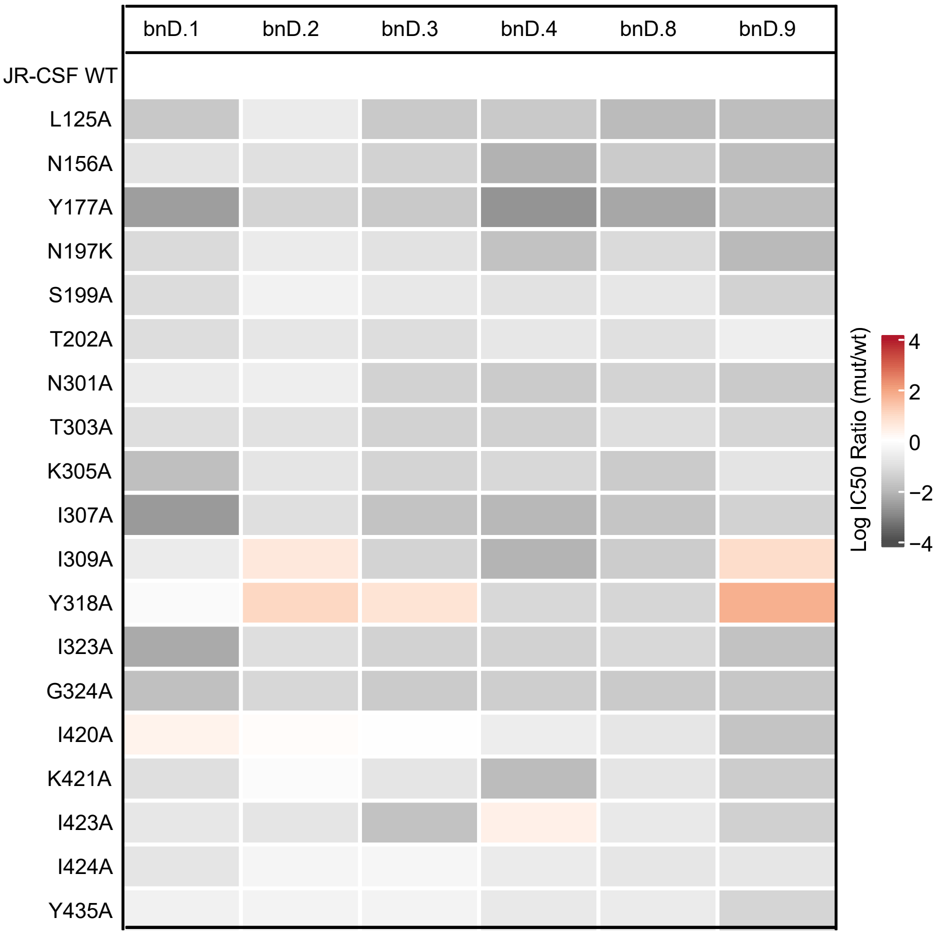

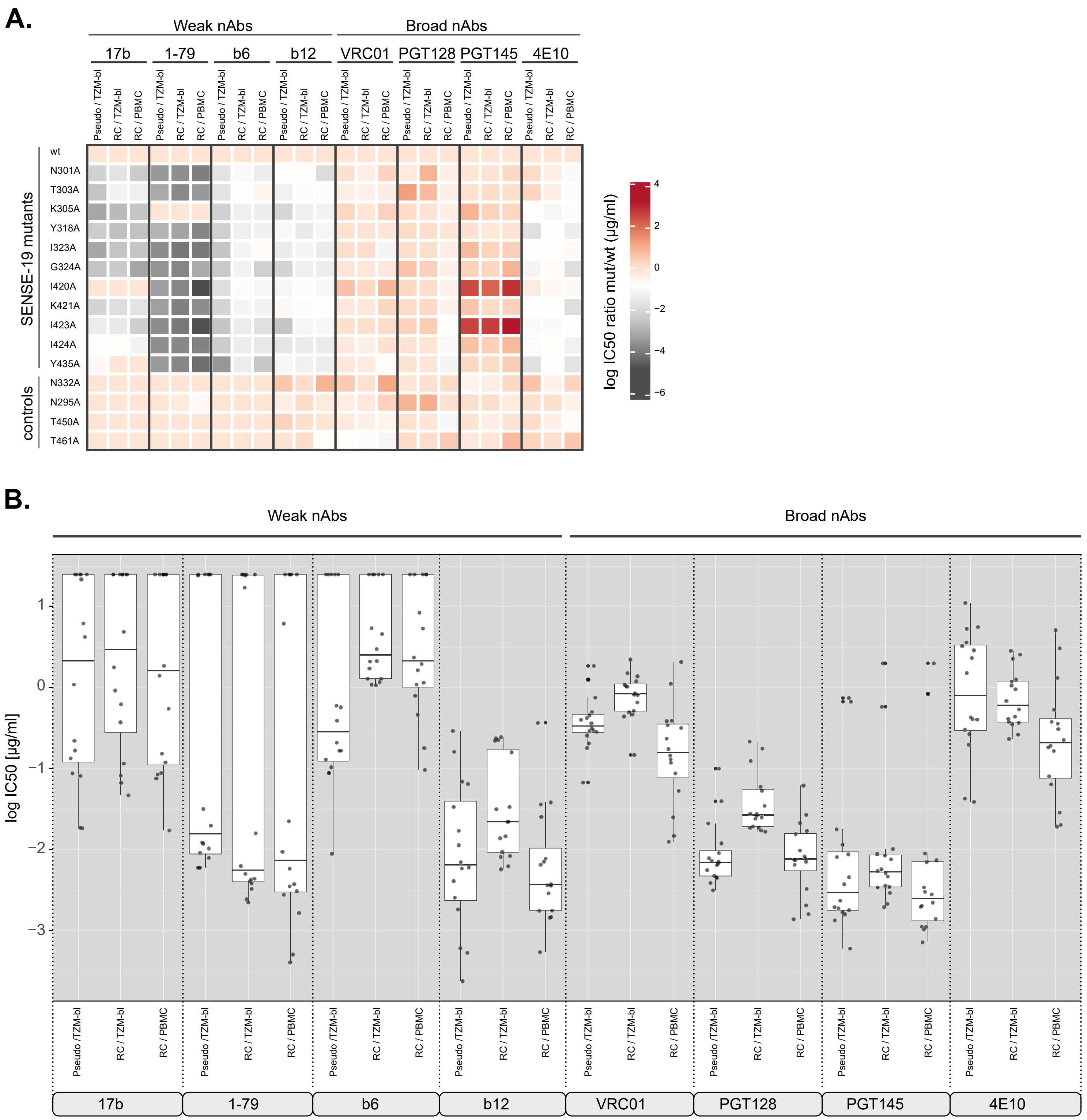

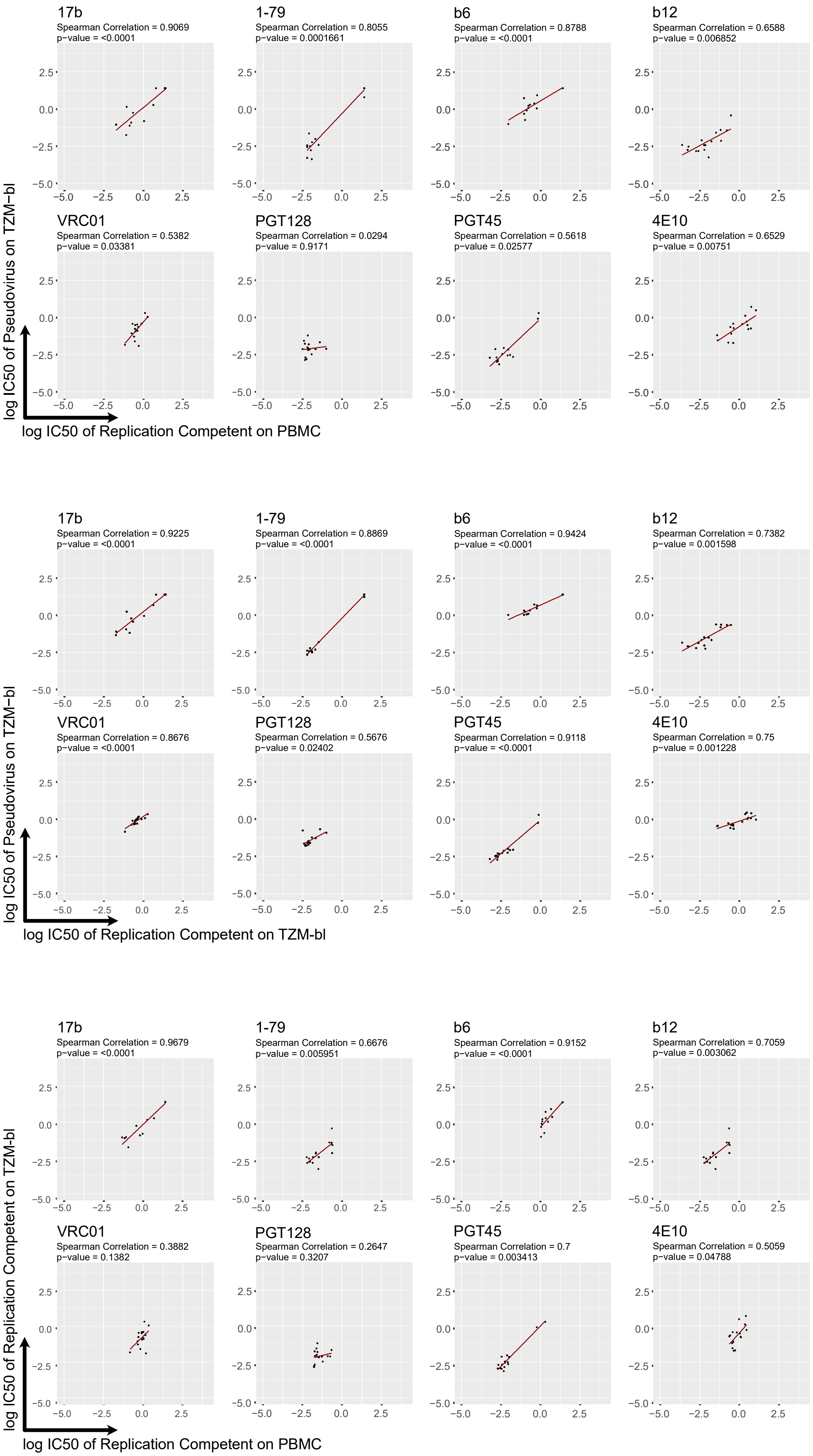

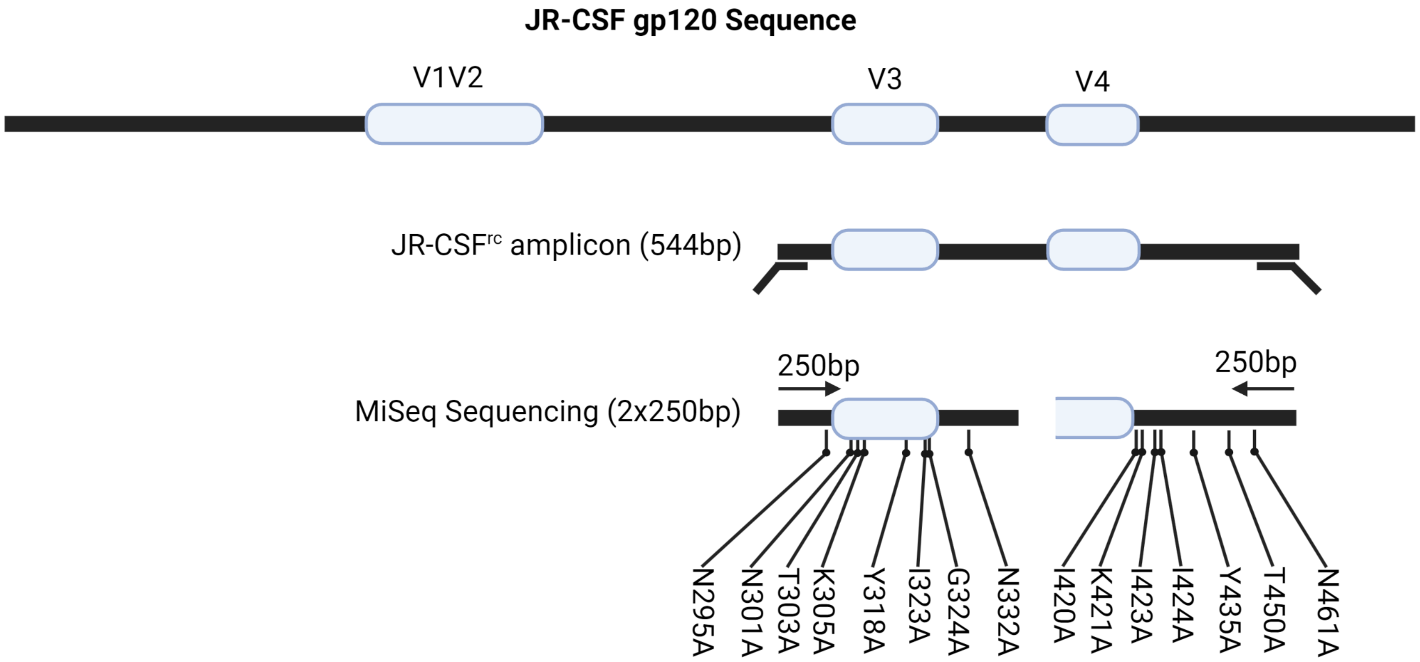
