## Supplemental Tables S1-S4 for "Assessing bnAb potency in the context of HIV-1 Envelope conformational plasticity"

**Table S1**

| Plasma | JR-CSF neutralizing dataset | JR-CSF non-neut | non-B data | VL | Subtype | CD4 | weeks after infection | weeks off ART |
|---|---|---|---|---|---|---|---|---|
| Z07-G3 | x | | | 19,000 | B | 313 | 304 | 193 |
| Z40-G1 | | x | | 33,400 | B | NA | 123 | 123 |
| Z36-G3 | x | | | 16,000 | B | 363 | 272 | 180 |
| Z92-G1 | | x | x | 8,600 | AE | 277 | 277 | 221 |
| Z38-G1 | x | | | 2,250 | B | 279 | 142 | 142 |
| Z39-G3 | x | | | 74,000 | B | 328 | 258 | 174 |
| Z91-G5 | x | | | 10,004 | B | 359 | 345 | 345 |
| Z49-G3 | x | | | 59,000 | B | 408 | 250 | 197 |
| Z48-G1 | x | | | 91,800 | B | 258 | 127 | 127 |
| Z62-G3 | | x | | 56,752 | B | 352 | 282 | 227 |
| Z75-G3 | | x | | 69,000 | B | 376 | 159 | 159 |
| Z71-G1 | | x | | 125,530 | B | 217 | 71 | 71 |
| Z02-G1 | x | | | 181,462 | B | 388 | 108 | 108 |
| S31438 | x | | x | 7,130 | A | NA | NA | NA |
| S51751 | | | x | 101,000 | C | NA | NA | NA |
| S31302 | x | | x | 54,500 | C | NA | NA | NA |
| S5138-G5 | | | x | 85,700 | AE | NA | NA | NA |
| S52611 | x | | | 1,551 | B | NA | 271 | 62 |

**Table S2**

| Name | Epitope | Reference | Source of expression plasmid | Source of protein |
|---|---|---|---|---|
| b12 | CD4-bs | Barbas III et al. 1992 PNAS. 89(19):9339-43 | D. Burton, The Scripps Research Institute, La Jolla, USA | H. Katinger and D. Katinger, Polymun, Vienna, Austria |
| b6 | CD4-bs | Barbas III et al. 1992 PNAS. 89(19):9339-43 | D. Burton, The Scripps Research Institute, La Jolla, USA | H. Katinger and D. Katinger, Polymun, Vienna, Austria |
| 1F7 | CD4-bs | Buchacher et al. 1994 AIDS Res Hum Retroviruses. 10(4):359-69 | | H. Katinger and D. Katinger, Polymun, Vienna, Austria |
| 3BNC117 | CD4-bs | Scheid et al. 2011 Science 16;333(6049):1633-7 | M. Nussenzweig, The Rockefeller University, New York, USA | Own production in 293T cells |
| PGV04 | CD4-bs | Wu et al. 2010 Science. 329(5993):856-61 | J. Mascola*, Vaccine Research Center, National Institutes of Health, Bethesda, USA | Own production in 293T cells |
| VRC01 | CD4-bs | Wu et al. 2010 Science. 329(5993):856-61 | J. Mascola*, Vaccine Research Center, National Institutes of Health, Bethesda, USA | Own production in 293T cells |
| NIH45-46 | CD4-bs | Scheid et al. 2011 Science 16;333(6049):1633-7 | M. Nussenzweig, The Rockefeller University, New York, USA | Own production in 293T cells |
| 1NC9 | CD4-bs | Scheid et al. 2011 Science 16;333(6049):1633-7 | Custom synthesized | Own production in 293T cells |
| CD4-IgG2 | CD4-bs | Allaway et al. 1995 AIDS Res Hum Retroviruses; 11(5):533-9. | | W. Olson*, Progenics Pharmaceuticals Inc |
| sCD4 | CD4-bs | Deen et al. 1988 Nature; 331(6151):82-4<br>Fisher et al. 1988 Nature;331(6151):76-8 | | W. Olson*, Progenics Pharmaceuticals Inc |
| sCD4-183 | CD4-bs | Ivan B. et al PLoS Biol. 2019; 17(1):e3000114. | Custom synthesized | Own production in E.coli |
| 17b | CD4i | Thali et al. 1993 J Virol. 67(7):3978-88 | Custom synthesized | J. Robinson*, Tulane University Medical Center, New Orleans, USA and own production in 293T cells |
| 2G12 | High Mannose Patch | Trkola et al. 1996 J Virol. 70(2):1100-8 | | H. Katinger and D. Katinger, Polymun, Vienna, Austria |
| PGT151 | Interface/Fusionpeptide | Falkowska et al. 2014 Immunity 40(5): 657–668. | D. Burton, The Scripps Research Institute, La Jolla, USA | Own production in 293T cells |
| 10E8 | MPER | Huang et al. 2012 Nature. 491(7424):406-12 | M. Connors*, National Institute of Allergy and Infectious Diseases, NIH, Bethesda, USA | Own production in 293T cells |
| 4E10 | MPER | Stiegler et al. 2001 AIDS Res Hum Retroviruses. 17(18):1757-65 | | H. Katinger and D. Katinger, Polymun, Vienna, Austria |
| 2F5 | MPER | Buchacher et al. 1994 AIDS Res Hum Retroviruses. 10(4):359-69 | | H. Katinger and D. Katinger, Polymun, Vienna, Austria |
| PGT145 | V2-Glycan | Walker et al. 2011 Nature. 477(7365):466-70 | D. Burton, The Scripps Research Institute, La Jolla, USA | Own production in 293T cells |
| PG9 | V2-Glycan | Walker et al. 2009 Science. 326(5950):285-9 | D. Burton, The Scripps Research Institute, La Jolla, USA | Own production in 293T cells |
| PG16 | V2-Glycan | Walker et al. 2009 Science. 326(5950):285-9 | D. Burton, The Scripps Research Institute, La Jolla, USA | Own production in 293T cells |
| PGDM1400 | V2-Glycan | Sok et al 2014 PNAS 111(49), 17624-17629 | D. Burton, The Scripps Research Institute, La Jolla, USA | Own production in 293T cells |
| PGT121 | V3 High Mannose Patch | Walker et al. 2011 Nature. 477(7365):466-70 | D. Burton, The Scripps Research Institute, La Jolla, USA | Own production in 293T cells |
| PGT128 | V3 High Mannose Patch | Walker et al. 2011 Nature. 477(7365):466-70 | D. Burton, The Scripps Research Institute, La Jolla, USA | Own production in 293T cells |
| PGT130 | V3 High Mannose Patch | Walker et al. 2011 Nature. 477(7365):466-70 | D. Burton, The Scripps Research Institute, La Jolla, USA | Own production in 293T cells |
| PGT135 | V3 High Mannose Patch | Walker et al. 2011 Nature. 477(7365):466-70 | D. Burton, The Scripps Research Institute, La Jolla, USA | Own production in 293T cells |
| BG18 | V3 High Mannose Patch | | Custom synthesized | Own production in 293T cells |
| 447-52D | V3-crown | Gorny et al. 1992 J Virol. 66(12):7538-42 | S. Zolla-Pazner (Icahn School of Medicine at Mount Sinai, New York, NY, USA) | H. Katinger and D. Katinger, Polymun, Vienna, Austria |
| 1-79 | V3-crown | Scheid et al. 2009 Nature. 458(7238):636-40 | M. Nussenzweig, The Rockefeller University, New York, USA | Own production in 293T cells |
| bnD.1 | V3-crown | Friedrich et al. 2021 Nat Commun. 2021; 12(1):6705. | Friedrich et al. 2021 Nat Commun. 2021; 12(1):6705. | Own production in E.coli |
| bnD.2 | V3-crown | Friedrich et al. 2021 Nat Commun. 2021; 12(1):6705. | Friedrich et al. 2021 Nat Commun. 2021; 12(1):6705. | Own production in E.coli |
| bnD.3 | V3-crown | Friedrich et al. 2021 Nat Commun. 2021; 12(1):6705. | Friedrich et al. 2021 Nat Commun. 2021; 12(1):6705. | Own production in E.coli |
| bnD.4 | V3-crown | Friedrich et al. 2021 Nat Commun. 2021; 12(1):6705. | Friedrich et al. 2021 Nat Commun. 2021; 12(1):6705. | Own production in E.coli |
| bnD.8 | αV3C | Glögl et al. 2023 Nat Struct Mol Biol. 2023; 30(9):1323-1336. | Glögl et al. 2023 Nat Struct Mol Biol. 2023; 30(9):1323-1336. | Own production in E.coli |
| bnD.9 | αV3C | Glögl et al. 2023 Nat Struct Mol Biol. 2023; 30(9):1323-1336. | Glögl et al. 2023 Nat Struct Mol Biol. 2023; 30(9):1323-1336. | Own production in E.coli |

* Through the NIH AIDS Reagent Program, Division of AIDS, NIAID, NIH

**Table S3:**

| clade | virus strain | Genbank entry | Tier |
|---|---|---|---|
| A | BG505_W6M_C2_T332N | DQ208458 | 2 |
| | KER2018.11 | AY736810 | 2 |
| | MG505.W0M.ENV.A2 | DQ208449 | 2 |
| | Q23_17 | AF004885 | 1B |
| | Q769_H5 | AF407159 | 2 |
| | Q842_D12 | AF407160 | 2 |
| B | ZEnv16_1202_7 | KU600818 | 2 |
| | ZEnv07_0504_15 | KU600814 | 2 |
| | ZEnv91_0505_12 | KU600815 | 2 |
| | BaL_26 | DQ318211 | 1B |
| | JR-CSF | AY669726 | 2 |
| | JR-FL | AY669728 | 2 |
| | NAB5pre_cl_1 | EU023923 | 2 |
| | NAB9pre_cl_106 | EU023928 | 2 |
| | PVO.04 | AY835444 | 2 |
| | QH0692, clone 42 | AY835439 | 2 |
| | REJO4541 clone 67 | AY835449 | 2 |
| | RHPA4259 clone 7 | AY835447 | 2 |
| | TRO clone 11 | AY835445 | 2 |
| | WITO4160 clone 33 | AY835451 | 2 |
| C | 25925_2_22 | EF117273 | 1B |
| | CAP45.2.00.G3 | DQ435682 | 2 |
| | CAP88_6mo.c10 | KU198436 | 2 |
| | Du156.12 | DQ411852 | 2 |
| | DU422.1 | AY043175 | 2 |
| | ZM106F.PB9 | AY424163 | 2 |
| | ZM214M.PL15 | DQ3885 | 2 |
| | ZM233M.PB6 | DQ388517 | 2 |
| | ZM249M.PL1 | DQ388514 | 2 |
| | ZM53_12 | AY423984 | 2 |
| G | T252_7 | EU513190 | 2 |
| | NAB13pre_cl_9 | EU023937 | 2 |
| | X2088_c9 | EU885764 | 2 |
| CRF01_AE | ZEnv32_0111_5 | KU600816 | 2 |
| | ZEnv92_1008_8 | KU600817 | 2 |
| | C1080.c3 | JN944660 | 2 |
| | CNE5 | HM215415 | 2 |
| | CNE59 | HM215422 | 2 |
| CRF02_AG | T250_4 | EU513189 | 2 |
| CRF07_BC | CNE40 | HM215414 | 1B |

**Table S4**

| PCR 1: Addition of Adapters | | | PCR 2: Barcoding | | |
|---|---|---|---|---|---|
| Reaction mix: | | Vol (µl) | Reaction mix: | | Vol (µl) |
| $H_2O$ | | 33 | molecular grade $H_2O$ | | 12.2 |
| 5x Kapa HiFi Buffer | | 10 | 5x Kapa HiFi Buffer | | 4 |
| dNTPs Kapa (10mM) | | 1.5 | dNTPs Kapa (10mM) | | 0.6 |
| Primer Fwd-JR-CSF (10µM) | | 1.5 | Index Primer 1 (10 µM) | | 1 |
| Primer Rev-JR-CSF (10µM) | | 1.5 | Index Primer 2 (10 µM) | | 1 |
| $MgCl_2$ Kapa (25mM) | | 0.5 | | | |
| Kapa HiFi enzyme | | 1 | Kapa HiFi enzyme | | 1 |
| DNA template | | 1 | DNA template | | 1 |
| PCR settings: | | | PCR settings: | | |
| Initial denaturation | 95°C, 3 min | | Initial denaturation | 95°C, 5 min | |
| Denaturation | 98°C, 20 sec | 30 cycles | Denaturation | 98°C, 20 sec | 17 cycles |
| Primer annealing | 65°C, 20 sec | | Primer annealing | 60°C, 15 sec | |
| Elongation | 72°C, 30 sec | | Elongation | 72°C, 30 sec | |
| Final elongation | 72°C, 5 min | | Final elongation | 72°C, 5 min | |
| Hold | 4°C | | Hold | 4°C | |
